# Allosteric Communication in the Gating Mechanism for Controlled Protein Degradation by the Bacterial ClpP Peptidase

**DOI:** 10.1101/2023.03.01.530711

**Authors:** Ashan Dayananda, T. S. Hayden Dennison, Hewafonsekage Yasan H.Fonseka, Mohammad S. Avestan, Qi Wang, Riina Tehver, George Stan

**Affiliations:** Department of Chemistry, University of Cincinnati, Cincinnati, OH 45221, United States; Department of Physics and Astronomy, Denison University, Granville, OH 43023, United States

## Abstract

Proteolysis is essential for the control of metabolic pathways and cell cycle. Bacterial caseinolytic proteases (Clp) use peptidase components, such as ClpP, to degrade defective substrate proteins and to regulate cellular levels of stress-response proteins. To ensure selective degradation, access to the proteolytic chamber of the double– ring ClpP tetradecamer is controlled by a critical gating mechanism of the two axial pores. Binding of conserved loops of the Clp ATPase component of the protease or small molecules, such as acyldepsipeptide (ADEP), at peripheral ClpP ring sites triggers axial pore opening through dramatic conformational transitions of flexible N–terminal loops between disordered conformations in the “closed” pore state and ordered hairpins in the “open” pore state. In this study, we probe the allosteric communication underlying these conformational changes by comparing residue-residue couplings in molecular dynamics simulations of each configuration. Both principal component and normal mode analyses highlight large-scale conformational changes in the N-terminal loop regions and smaller amplitude motions of the peptidase core. Community network analysis reveals a switch between intra- and inter-protomer coupling in the open - close pore transition. Allosteric pathways that connect the ADEP binding sites to N-terminal loops are rewired in this transition, with shorter network paths in the open pore configuration supporting stronger intra- and inter-ring coupling. Structural perturbations, either through removal of ADEP molecules or point mutations, alter the allosteric network to weaken the coupling.

## I. INTRODUCTION

Maintaining protein homeostasis at the cellular level is essential in all kingdoms of life^1,2^. Bacterial Caseinolytic proteases (Clp) assist these mechanisms by performing intracellular protein quality control through regulatory protein degradation^3^. Self-compartmentalized Clp nanomachines comprise a central barrel–like peptidase, such as ClpP, and one or two ring–shaped ATPase components, such as ClpA or ClpX, which are axially stacked at the two opposite ends of the peptidase^4,5^. Complex formation, which is dependent on ATP binding, tightly regulates the degradation process to prevent uncontrolled protein destruction^6,7^. The isolated ClpP is catalytically inactive as access to its proteolytic chamber is precluded by axial gates locked in a closed configuration, which allow diffusion of short peptides^8–10^ but hinder entry of longer unfolded polypeptide chains and block internalization of folded proteins^11–13^. Docking of one or two ATPase partners to ClpP triggers gate opening to unleash the powerful degradation mechanism^14–17^. Upon recognizing SPs targeted for degradation through short peptide tags attached covalently at one of the polypeptide terminals, the ATPase applies repetitive mechanical forces to effect SP unfolding and translocation through its narrow central channel and to propagate the unfolded polypeptide towards the peptidase core^18–25^. SP degradation takes place processively and yields small peptide fragments of 7-8 amino acids^26^.

Highly conserved N–terminal loops (amino acids 1-19) control access to the degradation chamber through the axial pores of the double-ring tetradecameric ClpP^27,28^. Intermediate “head” regions (residues 20-122 and 149-193) connect N-terminal loops to the inter–ring equatorial interface formed by interlocked “handle” regions (residues 123-148) of each protomer (Figure 1). In the “closed” pore configuration, N–terminal loops assemble into a mesh that involves strong inter–loop contacts and that occludes passage to the proteolytic chamber^29^. Removal of the N–terminal loops in ClpP variants abolishes the gating mechanism and enables even the isolated peptidase to indiscriminately destroy unfolded proteins. Functional control of degradation through complex formation with the ATPase is mediated by contacts formed with a ClpP ring at hydrophobic grooves located at peripheral sites from the axial pore. Although docking of the hexameric ClpA or ClpX ATPases to the heptameric ClpP rings involves a symmetry mismatch^1,27^, binding of six conserved ATPase loops, which contain the IGL or IGF sequence motif in ClpA or ClpX, respectively, ensures robust degradation activity^4,29–31^. Weaker complexes with only five IGL/F loops are functional, albeit degradation proceeds at a reduced rate^15^. The N–terminal loops are also involved in the complex formation through interaction with the ATPase pore–2 loops, however they form weaker contacts due to their greater conformational flexibility^30,32,33^. Remarkably, pore opening may be effected without assistance from the ATPase through strongly cooperative binding of seven acyldepsipeptide 1 (ADEP1) molecules to the peripheral ClpP sites (Figure 1)^2,34^. Kinetic studies indicate that ADEP1 establishes favorable interactions with the ClpP hydrophobic groove through its phenylalanine, β-methylproline and alanine moieties, and the aliphatic tail^35^. This triggers a dramatic conformational transition to a quasi–symmetric configuration of N–terminal β–hairpins pointing outward from the proteolytic chamber that results in nearly doubling the pore diameter^26,34,36,37^. Notably, the ClpP structure is very similar in the ADEP-bound and ClpX-bound configurations, with the global C_*α*_-based root-mean-square deviation of 0.8 Åin one ClpP ring and 0.6 Åin the pore region (defined by the seven N-terminal loops and adjacent helices)^5^. ClpP pore opening induced by ADEP binding yields powerful antibacterial action through uncontrolled destruction of bacterial proteins that is pursued for therapeutic applications against pathogenic *Staphylococcus aureus* and *Mycobacterium tuberculosis*^35,38–41^. Structural plasticity of ClpP probed in crystallographic and computational studies led to the hypothesis that transient exit channels form in the equatorial region to facilitate the release of peptide fragments resulting from the degradation process^2,42^.

**FIG. 1.**
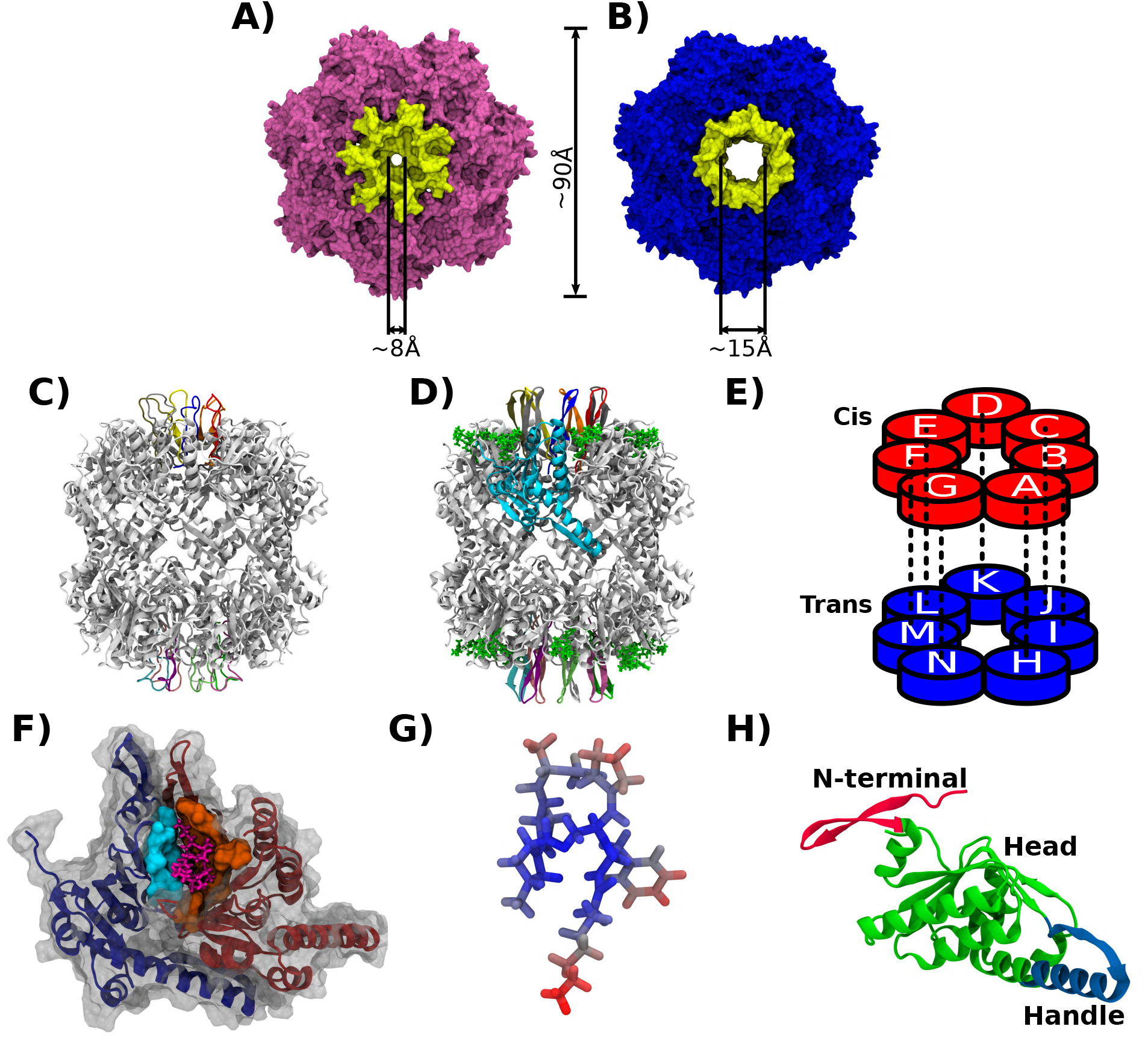
Structural details of the closed and open pore configurations of ClpP. (A)-(B) The crystal structure of the (A) closed (Protein Data Bank (PDB) ID: 1YG6) and (B) open (PDB ID: 3MT6) pore configuration of ClpP ((A) pink; (B) blue) reveals dramatic gate-controlling conformational transitions of N–terminal loops (yellow) (C)-(D) Side view of the (C) closed and (D) open pore configurations of ClpP (gray). N-terminal loops (color-coded), ADEP1 molecules (green) and one ClpP protomer (cyan) are highlighted. (E) Protomer organization of the cis (red) and trans (blue) heptameric rings and cross-ring inter-protomer coupling (dashes), mediated by the handle region interface, are indicated. (F) Top view of the ADEP binding site consisting of the hydrophobic pocket (indicated as a molecular surface) formed at the interface between chains A (blue) and B (orange). (G) ADEP1 molecular structure. (H) Domain architecture of ClpP monomer. N– terminal (residues 1-19, red), head (residues 20-122 and 149-193, green), handle (residues 123-148, blue) are highlighted. Molecular images in this work are rendered using Visual Molecular Dynamics^63^.

The details of allosteric communications that underlie the critical gate opening mechanism remain elusive even as multiple ClpP conformations have been resolved. Such challenges are commonly noted for ring-shaped molecular machines, such as GroEL, thermosome and CCT chaperonins, and are attributable to their large conformational plasticity, dynamic rewiring of allosteric networks and correlated intra- and inter-ring motions.^43–47^ To uncover the allosteric mechanisms in ClpP, we use computational approaches that are able to address such questions in diverse proteins by probing the complex networks of residue-residue interactions underlying the long-distance communication between the effector site and the functional site^48–51^. These approaches use concepts in graph theory combined with residue-residue coupling derived from structural data or conformational dynamics to identify “communities” of strongly coupled amino acids and to map the allosteric pathways connecting them. ^52–62^

In this paper, we describe comparative studies of ClpP conformational dynamics of ClpP in its open- and closed-pore configurations. To this end, we perform solvated, atomistic, molecular dynamics simulations of these systems and we identify the collective motions using principal component and normal mode analyses. Coupling between regions of the tetradecameric structure revealed by community network analysis indicate a switch between inter- and intra-protomer coupling as a result of the transition from the closed to the open pore configuration. Allosteric paths identified between the ADEP binding sites and the ClpP N-terminal loops highlight stronger intra- and inter-ring coupling between binding sites and N-terminal loops in the open configuration.

## II. METHODS

### A. Molecular Dynamics Simulations

The closed and open pore configurations of *Escherichia coli* ClpP were described using the X-ray structures with Protein Data Bank IDs 1YG6^29^ and 3MT6^34^, respectively. Unresolved regions of the X-ray structures were modeled using the ModLoop^64^ web server. To study the effect of perturbations on each structure, we considered several point mutations, which were implemented using PyMOL^65^ (Table S1). For each configuration and sequence molecular dynamics (MD) simulations were performed using the Gromacs^66^ 2022 package and the CHARMM36 all-atom force field^67^. The CGenFF server was used to generate force field parameters for the ADEP molecule compatible with the CHARMM36 force field^68^. Each protein structure was solvated in a dodecahedral box with dimensions ∼ 122 × 100 × 100 Å^3^ with water molecules represented using the CHARMM-modified TIP3P^69^ model. The system was neutralized by adding an appropriate number of Na ions for each setup (Table S1). The solvated system was energy minimized using the steepest descent algorithm for 50,000 steps with the convergence criterion of the maximum force value smaller than 1000 kJ/(mol· nm). Next, the systems were equilibrated by performing NVT and NPT dynamics.

First, simulations were performed for 500 ps in the NVT ensemble using the V-rescale^70^ algorithm, with T = 300 K, and by restraining the heavy atoms of the protein with a spring constant of 1000 kJ/(mol · nm^2^). In the second equilibration step, the restrained system was simulated for 500 ps in the NPT ensemble, with the constant pressure of 1 atm, using the Parrinello Rahman^71^ algorithm. The time step in all MD simulations was 2 fs. Finally, restraints were removed and, for each setup, four unbiased MD simulation trajectories (50 ns each) were performed in the NPT ensemble. For analysis purposes, data frames were saved every 100 ps were saved after excluding the first 10 ns of each trajectory. Root-mean-square deviations of simulations in each setup are shown in Figure S1.

### B. Data Analysis

### C. Dynamic Cross Correlation Matrix (DCCM)

To quantify the time-dependent directional correlations between residue pairs of the protein, we computed the Dynamic Cross Correlation Matrix (DCCM) of position fluctuations of C_*α*_ atoms of protein residues using the Bio3D package^72^. DCCM is an N × N matrix, with N equal to the number of residues, where each element C_*ij*_ corresponds to the dynamic cross-correlation between residues i and j:

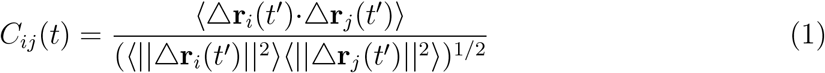

Here, r_*i*_ and r_*j*_ are the spatial positions of the C_*α*_ atoms of residues i and j as a function of time, ⟨·⟩ denotes the ensemble average over all trajectories and all time frames up to time t, and △r_*i*_(t^′^) = r_*i*_(t^′^) −⟨r_*i*_⟩ denotes the instantaneous position fluctuation of residue i from its mean over the simulation time. Correlation values range from -1 to 1, with positive C_*ij*_ values corresponding to motions of i and j atoms that are moving in the same direction, whereas the negative values indicate motions in opposite directions. Convergence of the DCCM matrix, shown in Figure S2, was assessed using R(t) = (1/N_*p*_) ∑_(*ij*)_ (C_*ij*_(t) − C_*ij*_(t − τ))^2^, where N_*p*_ is the number of residue pairs and the time interval τ = 5 ns^73^. We note that evaluation of R(t) using several τ values between 2.5 ns and 10 ns consistently yield R(t) ≤ 0.001. Here, C_*ij*_ is evaluated using data frames up to the total simulation time per trajectory t ⩽ 50 ns.

DCCM maps yield couplings between highly interconnected residue pairs that makes it challenging to decipher correlated motions of larger protein regions. To address this limitation, we performed community detection using the strongly coupled residue pairs in DCCM (|C_*ij*_| ⩾ 0.6) and the Girvan-Newman algorithm^74^ implemented in the cna() function in the Bio3D software package^72^. In this approach, the full residue network is split into highly intra-connected communities but weakly inter-connected between communities. In the Girvan-Newman algorithm, the number of communities is selected using the maximum modularity approach. Modularity quantifies how well a network is partitioned into communities. Since this is a probability measure, the values are in between 0 and 1. In general, for an ensemble, MD derived correlation network modularity above 0.7 indicates reasonable partitioning in a network^72,75^.

### D. Principal Component Analysis

Molecular Dynamics of complex biomolecular systems produce high-dimensional datasets comprising atomic positions saved in each time step. To glean information about the most significant conformational dynamics at a coarse-grained level one can employ Principal Component Analysis^76–78^. Here, we probe the functional dynamics between the open and closed configurations of ClpP by performing PCA. In PCA, the covariance matrix (⟨△r_*i*_·△r_*j*_⟩) comprising positional fluctuations is diagonalized to determine the set of eigenvalues and eigenvectors. PCA calculations are performed using the GROMACS analysis tools g covar and g anaeig^66^, where g covar generates both eigenvalues and eigenvectors by diagonalizing the covariance matrix and g anaeig filters the original trajectory and projects it along a given eigenvector. Prior to PCA calculation, we remove translation and rotation degrees of freedom of the entire molecule by superimposing each frame of the MD simulation onto the crystal structure. The comparison between the essential subspaces corresponding to open and closed configurations are performed by calculating the Root Mean Square Inner Product (RMSIP), which computes the overlap between two subsets of eigenvectors 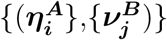 by using^79,80^

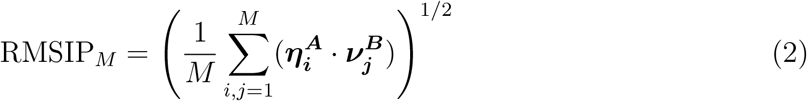

Here, the overlap is computed such that the top M eigenvalues in each configuration account for over 80% of the total variance.

### E. Optimal and suboptimal path analysis

To understand the dynamic coupling between the ADEP binding site and the N-terminal loop regions of the ClpP ring, we calculate the optimal and suboptimal paths traversing from each ADEP binding site to all the N-terminal loops in one heptameric ring. We use the cnapath() function implemented in the Bio3D package^58,72,75^. In the path analysis, each C_*α*_ atom is considered a node, and the connection between nodes is weighted by w_*ij*_ = −log(|C_*ij*_|). To remove weakly correlated regions and interactions between residues which are not in contact, the DCCM is filtered using the cuttoff value |C_*ij*_| ⩾ 0.3 and the dynamic contact map obtained in the MD trajectories. The contact map is generated in two steps. First, we identify persistent contacts in each trajectory, i.e. residue pairs with the C_*α*_-C_*α*_ distance d_*ij*_ ⩽ 10 Åpresent in at least 75% of the simulation time frames^59,81^. Next, the dynamic contact map is generated as the consensus map of contacts identified in at least three of the four trajectories. We find that the dynamic contact maps includes ≃ 84% of contacts present in the native structure and ≃ 2% new contacts. Paths are determined by using an efficient bidirectional approach that simultaneously initiates the search from one “source” (ADEP binding site) residue and one “sink” (N-terminal loop) residue and identifies closed paths upon locating common nodes. The path with the shortest length is identified as optimal, whereas slightly longer paths that are closed are identified as suboptimal. Accordingly, analysis of paths traversing between the “source” and the “sink” is useful to glean information about the allosteric regulation of regions that do not show large conformational changes. In our analysis, 300 paths are calculated for each “source” and “sink” residue pair and path length distributions are analyzed to assess the strength of the correlated motions. The extent of the overlap between two path length distributions, p_*i*_(x), i = 1, 2, is quantified by using the overlapping coefficient, OC = ⎰min{p_1_(x), p_2_(x)}dx.^82,83^ Further, the normalized node degeneracy, evaluated as the fraction of paths traversing through a given node, is used to identify the important residues that contribute to the allosteric network. Anaysis of node degeneracy indicates that approximately 350 paths between the source and sink are sufficient for convergence^59^.

### F. Normal mode analysis and Structural perturbation method

In order to calculate the normal modes of the proteins and analyze the response of the modes to perturbations, we modeled the proteins as elastic networks composed of N nodes where N is the number of amino acid residues in the structure^44,84,85^. The nodes are placed at the locations of the C_*α*_ atoms of the amino acid residues in the corresponding PDB structures. All nodes that are within a cutoff distance R_*c*_ = 9 Åare connected via harmonic potential with the energy function:

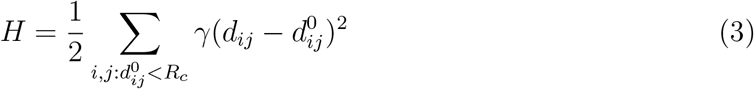

where γ is the spring constant that defines the energy scale, d_*ij*_ is the dynamic distance between residues i and j, and 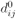 is the distance between the residues in the PDB structure. The normal mode calculation yields a set of 3N-dimensional eigenvectors,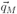 and corresponding eigenvalues ω_*M*_ for each mode M. We also analyze the responses of these modes to perturbations that mimic to point mutations of specific amino acids. This approach has been termed the Structural Perturbation Method (SPM) ^86,87^ and in practice we calculate the response to a mutation at the site i:

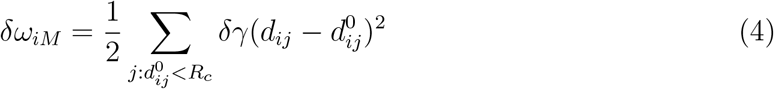

where δγ is the perturbed spring constant. The sum is over all other nodes in the network. The greater the response δω_*iM*_, the more dynamically significant a specific residue is to a given mode. We highlight the nodes that have δω_*iM*_ values that are three-fold above the average value for a mode as significant.

The overlap function quantifies how a given normal mode compares with the conformational change along a transition pathway. We compute the overlap function by projecting the eigenvector *q*_*M*_ of a given mode M onto the displacement vector between the open and closed configurations according to the formula^88^

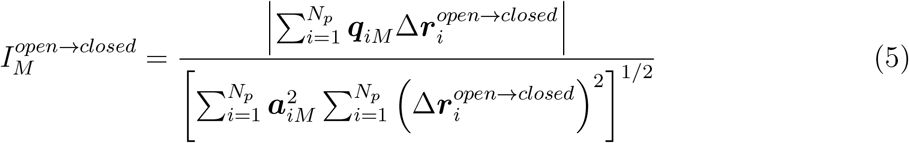

where the sum is over the *N*_*p*_ nodes, 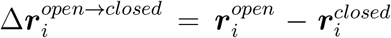, and 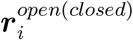 are position vectors of the ith node in the open (closed) structures, respectively. A value of one for the overlap corresponds to the direction given by the eigenvector *a*_*M*_ being identical with that of Δ*r*. The relative amplitude of node i in mode M is obtained using 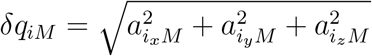 where 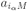, α = x, y, z, are the components of the normalized eigenvector *a*_*M*_.

## III. RESULTS

### A. Collective Motions Underlying the Gate-opening Conformational Transition Involve Primarily N-terminal Loop Regions of ClpP

To glean the collective motions of ClpP protomers underlying the gate-opening transition, we use Normal Mode Analysis (NMA) and Principal Component Analysis (PCA) (see the Methods section). These approaches have proved highly effective in studies of conformational changes and dynamics of multisubunit biomolecules, such as the ring-shaped chaperonins, ClpXP or immunoglobulin^45,89–92^. We use these analysis methods in conjunction to obtain detailed information about the collective motions, as inter-residue couplings in NMA are restricted to an elastic network model of the native protein structure, whereas in PCA they reflect dynamic, i.e. native as well as non-native, contacts. Nonlinear contributions to these couplings, quantified by considering the mutual information between residue coordinates^93^, have also been used to probe allosteric networks^58,60,61,94–97^. We note that a recent study of the tetracycline repressor dimer identified similar trends in the inter-residue coupling when comparing linear and nonlinear contributions^97^. A distinct advantage of networkbased approaches is their ability to probe the propagation of allosteric signal on timescales accessible to MD simulations, yet yielding results consistent with the much longer biological timescales^90,98^.

In PCA, diagonalization of the covariance matrix of atomic fluctuations yields the set of independent modes of motions of the protein that characterizes its essential subspace^77^. Eigenvectors of the matrix characterize the orthogonal directions of maximal variance, whereas eigenvalues determine the amplitude of positional deviations. We focus on the PC modes which correspond to the largest eigenvalues (Figure S5), which provide the major contribution to the variance of fluctuations. In the open state of ClpP, we find that eigenvalues corresponding to the top 3 PC modes contribute ≃ 40% of the cumulative variance of the fluctuations, and eigenvalues of the top 20 modes are required to explain ≃ 67% of the total variance. In the closed state, the top 3 eigenvalues contribute ≃ 52% of the total variance, whereas the top 20 eigenvalues contribute ≃ 74% of the total variance. In both cases, examination of the motions corresponding to the top 2 PC modes indicate a combination of large amplitude swing-type and torsional motions of the N-terminal loops that enable pore opening and closing (Figures 2A-B and Movies SM1-4) (Multimedia View). Handle domains in the inter-ring interface undergo slight contracting twisting motions. More specifically, in the open state, PC1 corresponds to swing-like motions that underlie pore opening and closing and PC2 corresponds to torsional motions of N-terminal loops. In the closed configuration, PC1 corresponds to torsional motions and PC2 corresponds to a combination of swing-like and torsional motions.

**FIG. 2.**
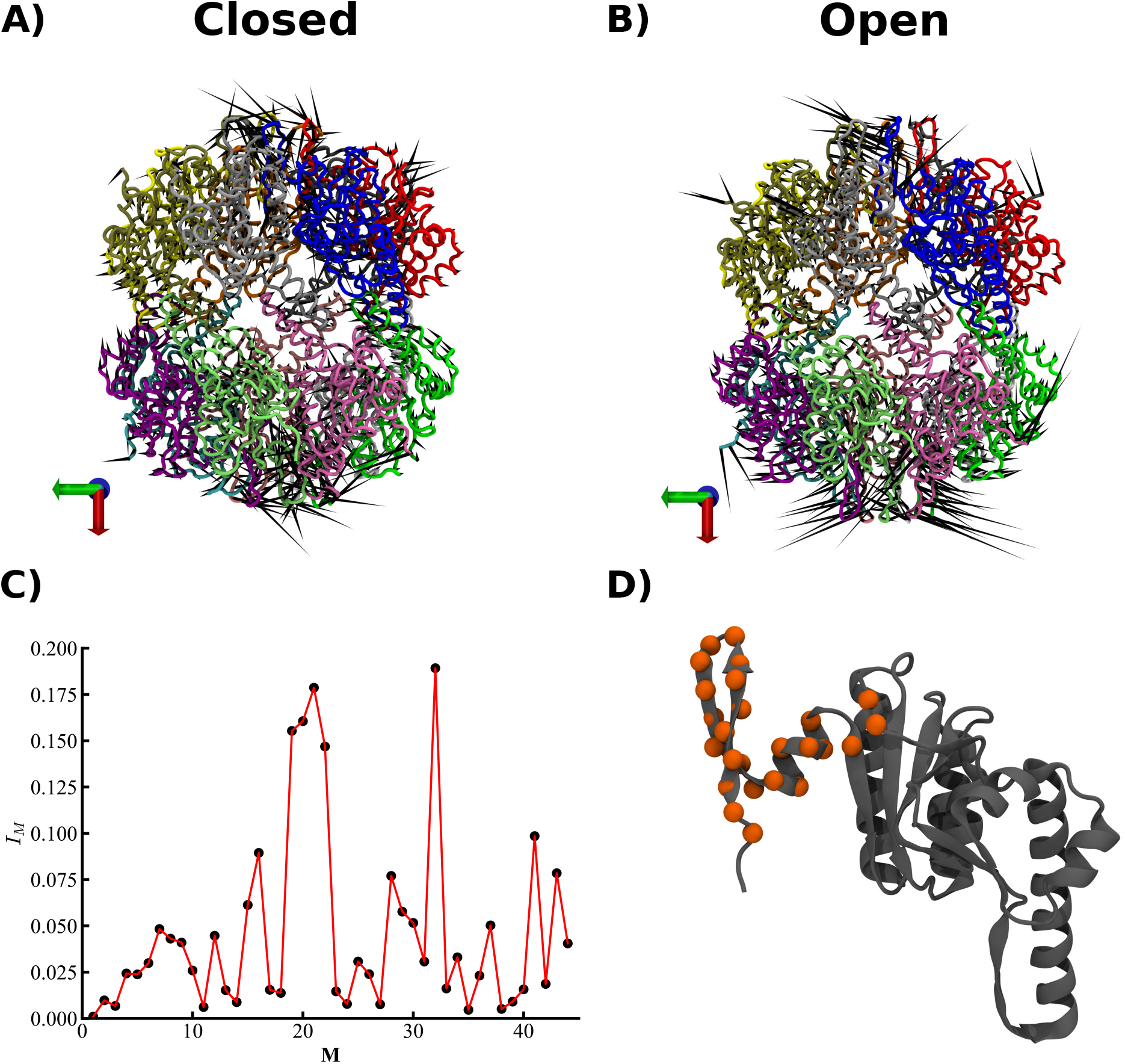
Principal Component and Normal Mode Analysis for ClpP conformations (A)-(B) Motions corresponding to the first principal component in (A) closed and (B) open pore configurations (Multimedia View). (C) Overlap of normal modes of ClpP with amino acid displacements in the open →− close transition. (D) The hot-spot residues (orange) extracted from the structural perturbation results. The list of hot-spot residues for the top two modes is shown in Table S2.

To discern the contribution of harmonic vibrations to these motions we perform normal mode analysis of the open state configuration (see Methods). We focus on the subset of normal modes whose eigenvectors have the largest overlap (see Methods) with the conformational changes corresponding to the transition between the open and closed conformations. As shown in Figure 2C, five modes have overlap 0.15 ≲ I_*M*_ ≲ 0.2, whereas all other modes have smaller contributions (I_*M*_ < 0.1). The absence of a single, dominant, mode indicates the lack of strong coordination of motions of the seven subunits of each ring to enable the transition. We also note that the lowest frequency modes, which describe global motions of the ClpP tetradecamer, have small contributions to the open → closed pore transition, whereas the five higher frequency modes, which describe more local motions, are more suitable to describe the transition. Consistently, the amplitudes of residue motions in the top five modes indicate large values only in the loop regions and negligible amplitudes outside of these regions (Figure S3). These results reveal that dominant conformational changes in this transition are primarily associated with motions of the N-terminal loops. On the basis of the comparison between PCA and NMA results, we surmise that dynamic fluctuations provide a stronger contribution in the handle and head regions than in the N-terminal loop regions.

Hot-spot residues that are critical for allosteric communication are identified by employing the Structural Perturbation method (see the Methods section). Here, the perturbations imposed by point mutations capture the change in the local network energy and hot-spot residues correspond to nodes whose response (δω_*iM*_) exceeds by three-fold the average value. Table S2 shows the hot-spot residues derived from the two modes with the largest overlap, 38 and 28 (Figure S4 and Movies SM5-6) (Multimedia View), and Figure 2D illustrates the location of these hot-spots residues projected onto a single ClpP monomer. We note that NMA and SPM analysis identify hot-spot residues clustered primarily in the N-terminal loop region, in accord with the dominant structural flexibility of this region in the harmonic approximation. In addition to the hot-spot residue cluster within and near the N-terminal loops, SPM analysis highlights two residues (His191, Arg192) in the C-terminal region.

Both PCA and NMA results are consistent with observations of structural studies, which highlighted that the peptidase core, ClpP(20-193), has virtually identical conformations in the open and closed configurations. On this basis, it was proposed that ADEP1 binding causes a significant conformational change in the N-terminal loops and only small-amplitude motions of the equatorial belt formed between two rings^34^. To quantify these contributions to the motions and to pinpoint the regions with the largest contribution to the open → closed transition, we further probe the PCs associated with the motions of the N-terminal loops and of the peptidase core, ClpP(20-193), respectively, in closed and open configurations. To this end, we perform separate PCA of each of these two ClpP regions in each of the two configurations and we determine the Root Mean Square Inner Product (RMSIP) of the two subsets of eigenvectors, which provides a measure of similarity of motions described by the PC modes (see Methods).

Comparison of collective motions of N-terminal loops was computed by considering the eigenvectors corresponding to the largest 11 eigenvalues, which collectively account for 80% of the variance, thus representing the essential subspace. Quantitatively, the overlap between the essential subspaces corresponding to N-terminal regions indicates weak similarity, with RMSIP ≈ 0.37. By contrast, for the peptidase core, ClpP(20-193), where the top 110 eigenvectors must be included to describe the essential subspaces, we find a strong overlap between the PC modes, with RMSIP ≈ 0.73. This indicates that ADEP binding has only a weak effect on the dynamics of the peptidase core.

Overall, NMA and PCA data suggest that the conformational transition of ClpP protomers during the gate opening and closing are best described by an ensemble of modes. Taken together, both PC and normal mode data reveal that large conformation changes only exert at the N-terminal loop regions while the core remains mainly intact.

### B. Distinct Coupling between ClpP regions in Closed and Open Pore Configurations

To investigate the correlated motions of regions in the ClpP tetradecamer, we employ the community network analysis, which uses the residue-residue coupling quantified by the directional cross-correlation map (DCCM, see the Methods section). DCCM maps are highly interconnected at the residue level, which makes it complicated to extract information for large systems. Therefore, we probe community network clusters by converting the atomic cross-correlations to a coarse-grained type community network using the Girvan-Newman clustering method^74^. To investigate both intra- and inter-protomer coupling, we select the cutoff of the cross-correlation between residues i and j, |C_*ij*_|, such that the maximum modularity (see Methods and Figure S6) corresponds to a larger number of community clusters than the number of ClpP protomers, 14, in each of the three ClpP configurations. We find that this requirement is satisfied by |C_*ij*_| ⩾ 0.6, which yields ≈ 30 − 40 community clusters in ClpP configurations (Tables S3-S5). As shown in Figure 3, the community network analysis reveals distinct patterns of domain coupling in ADEP-bound and unbound conformations of ClpP. Strikingly, the ADEP-bound open conformation is characterized by extensive intra-subunit coupling within six subunits that involves strong coordination between the N-terminal loop, handle and head regions (Figure 3A). By contrast, limited coupling is observed across the equatorial interface between protomers of the cis (A-G) and trans (H-N) rings (according to the ClpP protomer organization shown in Figure 1), involving the handle regions of two protomer pairs (C-J and D-K). Removal of ADEP from the open pore conformation results in dramatic changes in intra-protomer coupling, which nearly abolish correlation between the handle region and the loop and head regions (Figure 3B). Furthermore, the inter-ring coupling is enhanced through coordinated motions of handle regions of five protomer pairs. These changes yield coupling patterns similar to the closed conformation, in which intra-protomer coupling is weaker and inter-ring coupling between handle regions of protomer pairs is prevalent (Figure 3C). We note that these results are consistent with the important role of the handle domain in tetradecamer formation and stacking of the two rings (cis and trans)^99^.

**FIG. 3.**
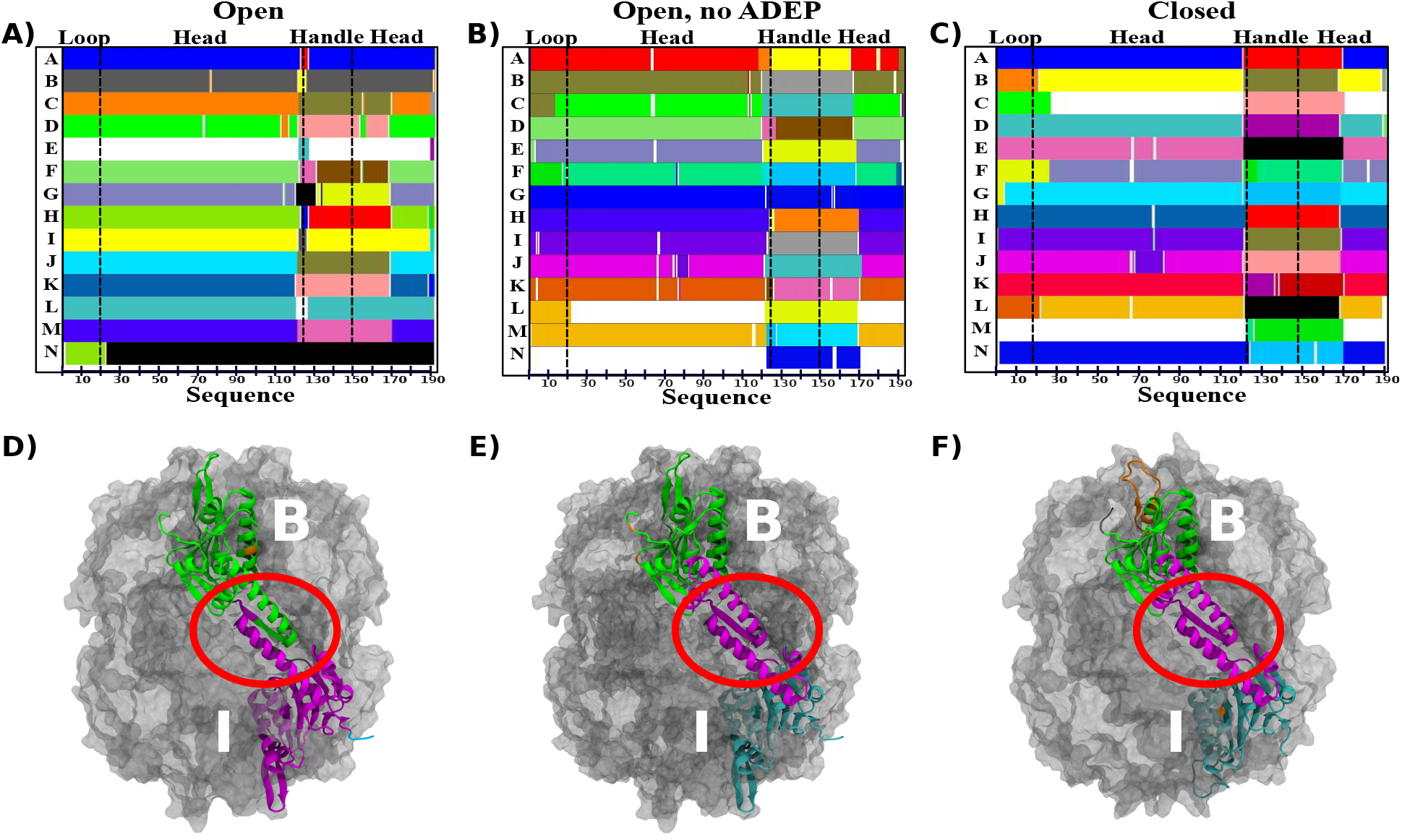
Community network analysis for ClpP tetradecamer configurations. Community maps for (A) open pore (B) open, no ADEP bound (C) closed pore configurations. The N-terminal loop, head, and handle regions are indicated for the cis (protomers A-G) and trans (H-N) rings of ClpP. (D)-(F) Structural details of inter-protomer coupling across the ClpP rings. (D) Strong intraprotomer and weak inter-protomer coupling in the open pore configuration (E)-(F) Strong interprotomer coupling mediated by the equatorial interface, including the handle region (highlighted by red elipse), in (E) open, no ADEP and (F) closed pore configuration. Communities identified in (A)-(C) are highlighted.

### C. Optimal and Suboptimal Path Analysis Reveal Stronger Coupling between Allosteric and Active Sites of ADEP bound ClpP

The absence of large-scale rigid-body domain motions of ClpP protomers and the proximity of the ADEP binding sites to the N-terminal loops suggest that allosteric signals can be transmitted through relatively short, intra-protomer, paths. The ring structure of ClpP, however, also allows effective allosteric communication to take place between ADEP binding site of one protomer and N-terminal loops of the other intra-ring protomers. To probe these allosteric mechanisms in a quantitative manner, we use approaches that combine residueresidue positional correlations and concepts in graph theory (see Methods). To this end, we map the allosteric paths that connect one ADEP binding site, termed “source”, and the seven N-terminal loops within one ring, or “sink”. In the correlation network, each node represents one protein residue and the connecting edges have associated lengths that reflect the cross-correlation values (see Methods)^54^. The path length between nodes located in the source and sink is then identified as the sum of the lengths of all individual edges that connect these nodes. We emphasize that, given the construction of allosteric map in the correlation space, the relative importance of the allosteric paths depends on the strength of the coupling between constituent residue pairs rather than their proximity in the physical space. In this framework, the shortest paths between residues of the source and the sink reveal the strongest allosteric couplings within a protein^51^. Interestingly, the minimal length, or “optimal”, path was not found to be very sensitive to changes in the protein configuration, therefore analysis of allosteric communication limited to this path may yield misleading conclusions about the signaling pathways^59^. Instead, a comprehensive analysis requires the additional evaluation of “suboptimal” paths, which have slightly longer lengths than the optimal path^59^.

Using this framework, we computed 300 paths for each residue pair formed by one source and one sink amino acid. To this end, we select as source one ADEP binding site (Figure 4A), comprising residues Val44, Leu48, Phe49, Glu51, Ala52, Phe82 from protomer i − 1, and Arg22, Leu23, Val28, Phe30, Tyr60, Tyr62, Ile90, Met92, Leu114, Leu189 from the adjacent protomer i, and as sink two representative residues, Thr10 and Gly13, located near the turn of each N-terminal loop in the same (cis) ring. The selection of representative loop residues suffices for the purposes of mapping paths, as allosteric signal propagates within the loop exclusively through intra-loop residues. By considering paths connecting each ADEP binding site to each loop, we examined a total of 470400 pathways traversing one ClpP ring (see Methods). In all three ClpP configurations, the optimal pathways are intra-protomeric and connect Arg22 to loop residues 13-17 (Table S6). Thus, optimal pathways are largely stable among the three ClpP configuration, which is in accord with observations made on small proteins noted above.

**FIG. 4.**
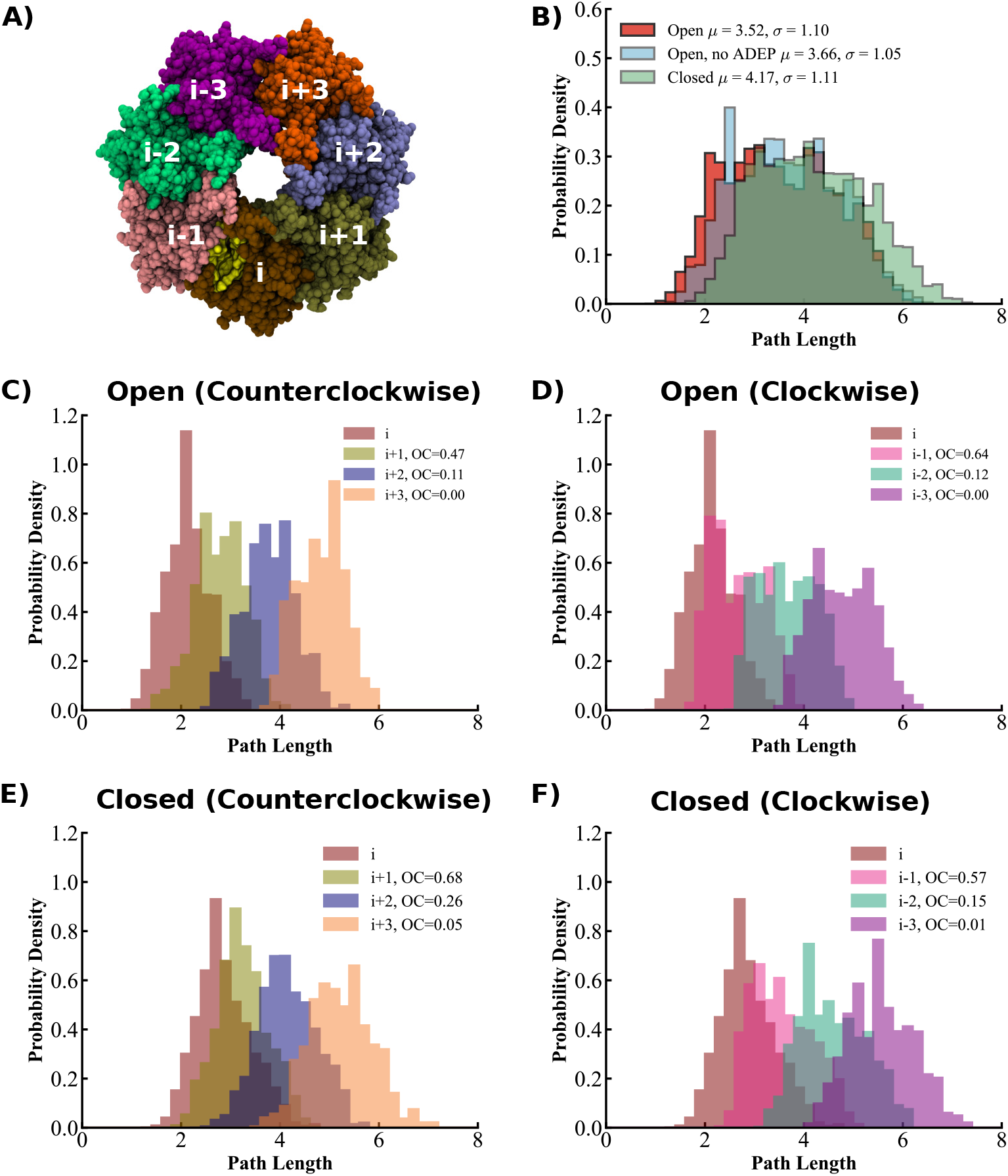
Path length distributions in allosteric signaling in ClpP configurations. (A) Intra-ring protomer ordering relative to a given ADEP binding site, *i* (yellow). Allosteric paths are mapped between the binding site and the N-terminal loops in the same protomer *i*, and in protomers in counterclockwise 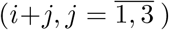and in clockwise 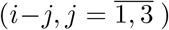 directions. (B) Probability density distributions of path lengths are shown for the complete set of paths between each binding site and all intra-ring loops in the open (red); open, no ADEP (blue); and closed (green) configurations of ClpP. The legend indicates mean and standard deviations of distributions. (C)-(D) Path length distributions of paths mapped in the open ClpP configuration between binding site *i* and loops in protomers in (C) counterclockwise and (D) clockwise directions are compared with the path length distribution of paths to the loop in protomer *i*. (E)-(F) Same as in (C)-(D) for the closed ClpP configuration. The legends indicate the overlapping coefficient, OC, between each distribution and the same protomer distribution.

In order to characterize the strength of allosteric communication from a broad perspective, we examine the set of suboptimal paths in each pore configuration as well as the subsets of paths connecting each ADEP binding site to a specific loop (Figure 4A). As shown in Figure 4B, the histograms of suboptimal path lengths in the three ClpP configurations highlight the stronger coupling between binding sites and loops effected by ADEP binding, with the shortest paths corresponding to the open state. By contrast, in the closed state, the path length distribution is shifted towards longer paths, with lengths up to ≃ 8, which indicates a weaker coupling between the ADEP binding site and the N-terminal loops. Perturbation of the open configuration through removal of the ADEP molecules results in a slight shift of the suboptimal path length distribution towards paths of intermediate lengths between those found in the open and closed configurations. Next, we probed the intra-ring propagation of the allosteric signal by examining the path length distributions corresponding to paths connecting each binding site to the N-terminal loop of each protomer (Figures 4C-F). In accord with the above observations, path lengths corresponding to the open state are consistently shorter than those for the closed state. In both open and closed pore configurations, the shortest paths originating from the ADEP binding site of protomer i correspond to paths to the N-terminal loop of the same protomer (Figures 4C-F). In the open state, we find relatively short paths, of length ≃ 2, to the nearest neighbor loops in both clockwise (CW) and counterclockwise (CCW) directions. This strong intra-ring coupling supports the ability of the hexameric ATPase to trigger ClpP gate-opening even with substoichiometric occupation of distal binding sites. Decreasing coupling strength is found to the N-terminal loops of the second and third nearest neighbor protomers, however, with slightly shorter paths in the CW direction (Figures S7A-C). In the closed state, we note the larger overlap than in the open state between path length distributions corresponding to neighboring loops and that of the same protomer loop, which indicates a slower intra-ring decay of the weaker allosteric signal in this state. In addition, the CW and CCW distributions corresponding to N-terminal loops of equidistant nearest neighbors overlap nearly completely (Figures S7D-F).

The distinct pattern of path lengths in the open and closed configurations of ClpP, as well as the marked effect of perturbations on the path lengths indicate a significant dynamic rewiring of allosteric communication. To obtain the microscopic understanding of the changes induced by perturbations, we probe the detailed paths in each configuration. To this end, we examine the three-dimensional maps associated with suboptimal paths (Figure 5). The structural maps of allosteric pathways in the open configuration indicate strong signaling propagated from the ADEP binding site to the nearest three N-terminal loops in the cis ring (Figure 5A). Removal of the ADEP molecules yields weaker coupling (indicated by thinner lines connecting the nodes) and increasing number of paths connecting the ADEP binding site to the more distant loops (Figure 5B). In the closed configuration, paths that connect multiple loops are increasingly found (Figure 5C).

**FIG. 5.**
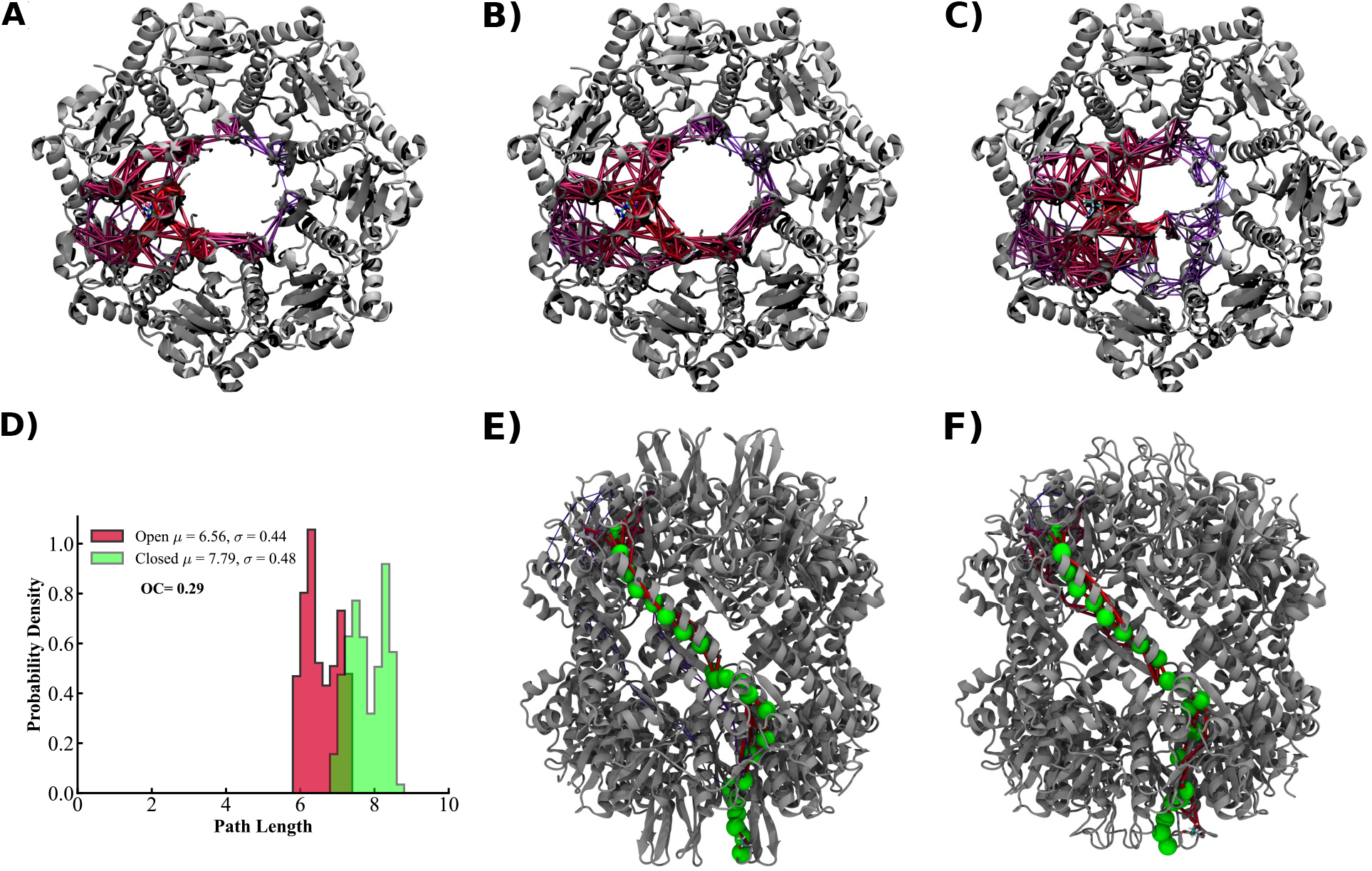
Signaling pathways for ClpP configurations derived from the dynamic network analysis. (A)-(C) Suboptimal paths (red) between one ADEP binding site and all N-terminal loops in the same ClpP ring (gray) in (A) open; (B) open, no ADEP; and (C) closed pore configurations. (D) Probability density distributions of suboptimal inter-ring paths in open (red) and closed (green) pore configurations that connect the ADEP binding site of a cis ring protomer and the N-terminal loop in its trans ring protomer partner. The mean and standard deviations of distributions and the overlapping coefficient between them are indicated. (E)-(F) Structural details of the optimal inter-ring paths in (E) open and (F) closed pore configurations.

Interestingly, as shown in Figure 5D, the analysis of inter-ring pathways connecting the binding site of one protomer (B) in the cis ring and the N-terminal loops of its partner (I) in the trans ring, using |C_*ij*_| ⩾ 0.6, reveals shorter paths, and therefore tighter inter-ring coupling, in the open configuration compared with the closed configuration. In the open configuration, allosteric communication between rings is primarily mediated by paths with lengths ⩽ 6.5, which are not available in the closed configuration, and the overlap between the two distributions is small, with the overlapping coefficient OC ≃ 0.29 (Figure 5D). Although, as noted in the previous section, the inter-ring handle interface is abolished in the open configuration, shorter paths become available through alternate routes, as indicated in Figures 5E-F. The shorter path lengths corresponding to the open configuration can be rationalized in terms of the connectivity between community networks of the cis and trans protomers that mediate the inter-ring allosteric communication, indicated in Figure 3. Whereas, in the closed configuration, paths must cross two gaps between intra-protomer communities to connect with the inter-protomer handle community, in the open configuration, a single, inter-protomer, gap must be crossed to connect the cis and trans protomer communities (Figures 3 and 5E-F). The differential gap penalty results in a length of ≃ 5.9 for the optimal path of in the open configuration, and ≃ 6.9 in the closed pore configuration, even as the structural details of the paths are similar (Table S7). We surmise that the handle interface, rather than facilitating inter-ring allosteric communication, acts as a constraint in the closed configuration, resulting in two intra-protomer path length penalties. By contrast, in the open configuration, the inter-protomer constraint is removed and a single path length penalty is applied.

Next, to identify residues that are critical for allosteric signaling, we compute the normalized intra-ring node degeneracy, i.e. the fraction of paths that include each node (see Methods). Table I summarizes the residues in the cis ring that have the node degeneracy ⩾ 0.10 in at least one configuration, as revealed by path analysis. The structural location of residues that act as critical nodes is highlighted in (Figure 6). Notably, residues Ile19_*i*_, Tyr20_*i*_, Ser21_*i*_, Ser21_*i*−1_, and Leu24_*i*−1_, at the base of N-terminal loops proximal to the ADEP binding site and Gln46_*i*−1_ and Leu50_*i*−1_, which are in the proximity of ADEP binding site, have node degeneracy ⩾ 0.10 in all three setups and therefore are likely to have a critical contribution to allosteric communication. (In our notation, residue labeling includes as subscript the protomer location in the cis ring, with the “source” binding site occupying protomers i − 1 and i. Protomers are numbered in the counterclockwise direction in the top view of the ring.) One set of residues, Ile19_*i*_, Ser21_*i*−1_, Leu24_*i*−1_, and Leu50_*i*−1_, has a slightly reduced degeneracy in the closed pore configuration and upon removal of ADEP molecules in the open configuration, whereas the other residues have increased degeneracies.

**TABLE 1.** Normalized node degeneracy derived from suboptimal path analysis. Residues are grouped according to the ClpP protomer they belong to (shown as a subscript). Protomers are numbered in the counterclockwise direction, in the top view of the cis ring. In the path analysis, the ADEP binding site “source” includes residues from protomers i and i-1. Highlighted residues represent critical nodes (degeneracy ⩾ 0.10) in all three ClpP configurations examined.

| Residue | Ser21<sub>$i-1$</sub> | Leu24<sub>$i-1$</sub> | Ala45<sub>$i-1$</sub> | Gln46<sub>$i-1$</sub> | Leu50<sub>$i-1$</sub> |  |
| --- | --- | --- | --- | --- | --- | --- |
| Open | **0.17** | **0.26** | 0.07 | **0.14** | **0.19** |  |
| Open, no ADEP | **0.17** | **0.26** | 0.08 | **0.14** | **0.15** |  |
| Closed | **0.14** | **0.17** | 0.11 | **0.14** | **0.17** |  |
| Residue | Ile19<sub>$i$</sub> | Tyr20<sub>$i$</sub> | Ser21<sub>$i$</sub> | Leu24<sub>$i$</sub> | Ile29<sub>$i$</sub> | Ser21<sub>$i+1$</sub> |
| Open | **0.22** | **0.11** | **0.19** | 0.11 | 0.10 | 0.06 |
| Open, no ADEP | **0.11** | **0.19** | **0.26** | 0.12 | 0.08 | 0.13 |
| Closed | **0.16** | **0.23** | **0.22** | 0.05 | 0.09 | 0.06 |

**FIG. 6.**
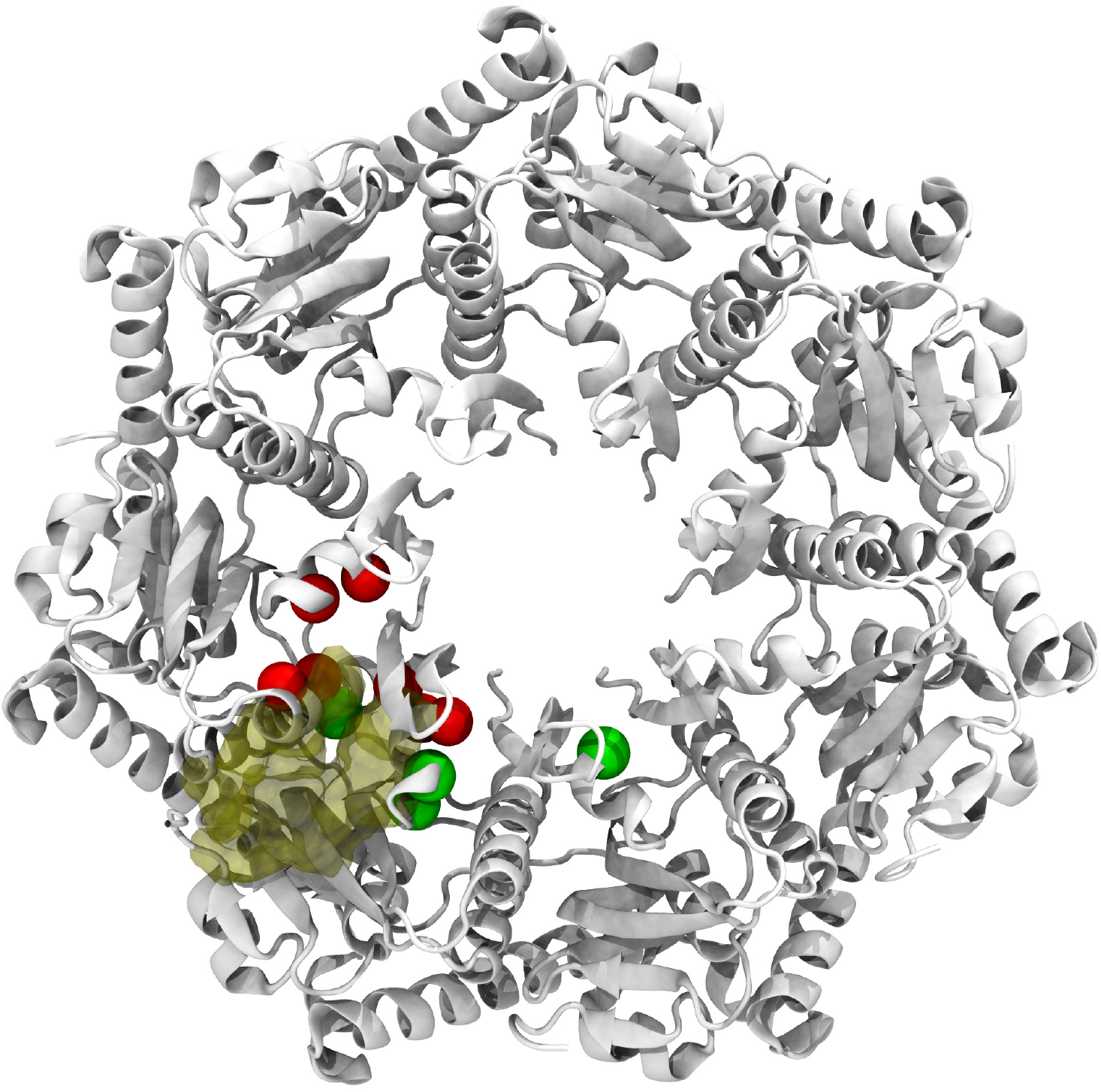
Hot-spot residues for ClpP configurations derived from the dynamic network analysis. Residues with similar (distinct) node degeneracy in open and closed pore configurations are indicated with red (green) dots.

Interestingly, we also note that residue Gln46_*i*−1_ shows no change in node degeneracy values in the three setups, indicating a weak sensitivity to structural perturbations. Structural studies revealed that the presence of Ile19 is crucial for the stability of the N-terminal loops and substrate translocation in the ADEP bound open state^100^. We note that Ile19 is present in all 3 setups with a slightly higher value in the open setup.

### D. N-terminal Mutations Differentially Alter Allosteric Communication in Open and Closed Pore Configurations

We further explore how perturbations alter the coupling between the allosteric and active sites of ClpP by engineering point mutations at N-terminal sites indicated to be functionally important (see Methods and Table S1)^100^. We focus, on the one hand, on single point-mutations, such as I7P, E8K, and K25E, that alter the stability of individual N-terminal loops, and, on the other hand, on double, E14A-R15A, and triple, E14A-R15A-E8A and E14A-R15A-K25A, mutations that affect intra- as well as inter-loop stabilization (Figure 7). To this end, we compare the path length distributions corresponding to shortest 5000 suboptimal paths among the allosteric pathways connecting one binding site to all the ClpP N-terminal loops in the cis ring (see Methods). As shown in Figure 7, we find that mutations have distinct effects on allosteric coupling, which can manifest differently in the open and closed pore configurations. Single mutations have generally large effects in both open and closed configurations. As shown in Figure 7B, the I7P mutation, which makes the coil conformation more favorable, has a drastic effect on the open configuration, with a low OC ≃ 0.22, in accord with the deleterious effect of this mutation on polypeptide degradation^100^. E8K and K25E mutations, which remove the stabilizing salt bridge at the base of the N-terminal loop, affect both the open and closed configurations (Figure 7B-C, which is consistent with their diminished degradation rates of both polypeptides, which stringently require an open gate, and peptides, which may be internalized through the closed gate^100^. Double and triple mutations including the E14A-R15A mutations that provide stabilizing inter-loop salt bridges, have a lesser effect on the closed than on the open pore configuration, as the salt bridges formed by Glu14 and Arg15 residues in neighboring protomers are not present in the closed state even in the wild-type ClpP (Figures 7D-E). We find that the triple mutation E14A-R15A-K25A has the strongest effect on allosteric paths, with OC ≃ 0.14 in the open configuration, which is in accord with the largest reduction in the degradation rate of polypeptides compared with the single and double mutations^100^.

**FIG. 7.**
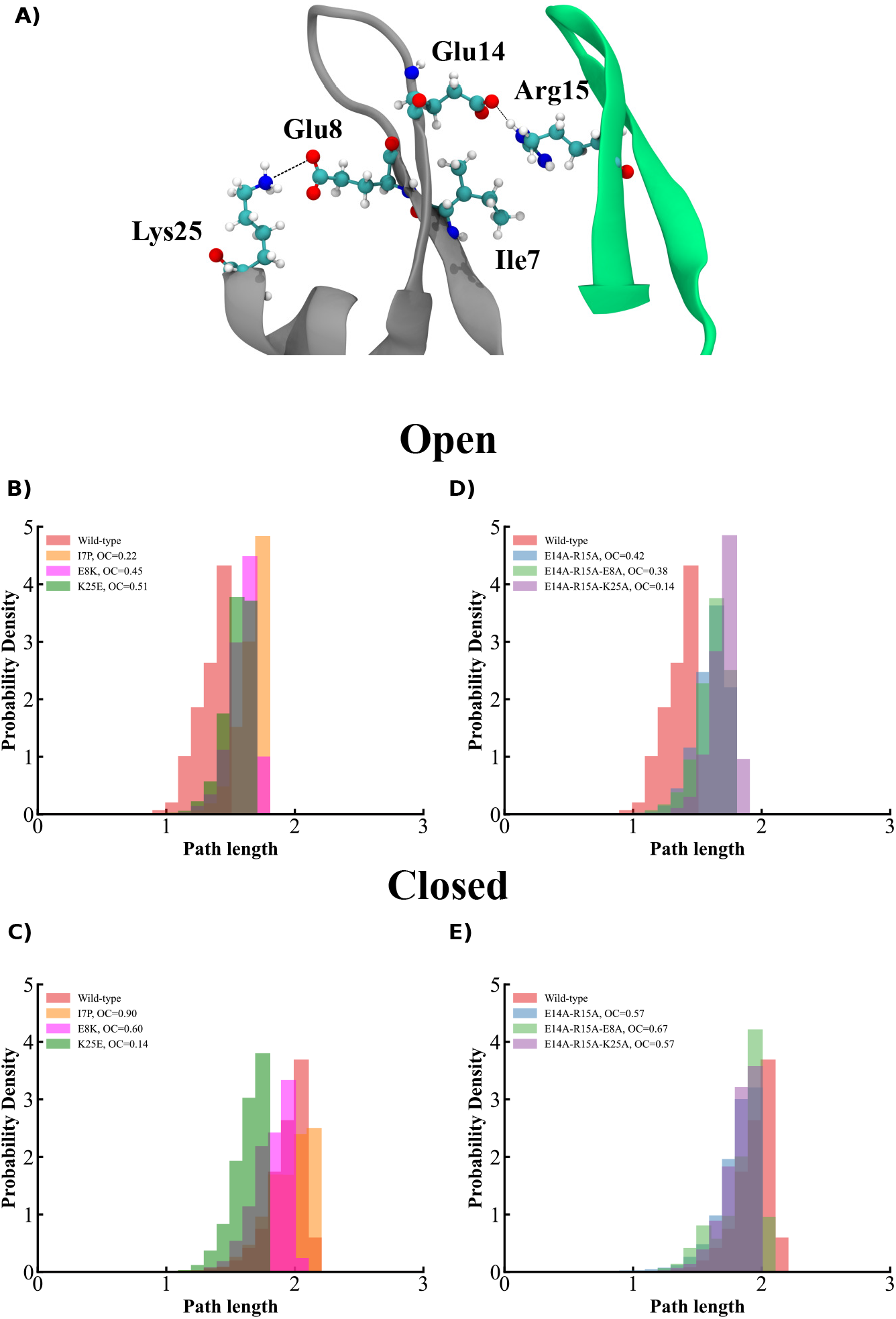
Effect of mutations on allosteric signaling. (A) Single point-mutations considered, I7P, E8K, and K25E, affect stability of individual loops (gray) and double and triple mutations, E14A-R15A, E14A-R15A-E8A, and E14A-R15A-K25A, affect both intra-loop and inter-loop (gray and green) stabilization. Salt bridges formed in the open pore configuration are highlighted. (B)-(C) Comparative probability distribution of the shortest 5000 paths in wild-type and single mutants in (B) open and (C) closed pore configurations. (D)-(E) Same as in (B)-(C) for double and triple mutants. Overlapping coefficients between each mutant and the wild-type distribution are indicated in the legend.

## IV. DISCUSSION

Our computational studies probe the allosteric mechanisms of the ClpP peptidase in response to effectors that activate its gate-opening conformational transition. Intriguingly, this transition involves limited structural rearrangement outside of the gate-controlling N-terminal loop region. How is the allosteric signal propagated in the absence of large-scale rigid-body motions of ClpP subunits? To address this question, we undertook comparative studies of the open and closed pore configurations of ClpP by performing equilibrium MD simulations of each of these states. Our results, quantified through principal component and normal mode analysis, highlight the similarity of motions of the peptidase core in the two states even as the loop motions are significantly different.

In our analysis, both structural perturbation, derived from Normal Mode calculations, and node degeneracies, computed using the positional cross-correlations, highlighted a set of hot-spot residues that are critical for the gate opening transition of ClpP. We find that the hot-spots derived from the harmonic approximation used in NMA reveal regions that are highly flexible and dynamic through their proximity to the N-terminal loop regions or the C-terminal regions. Node degeneracy values derived from dynamic cross-correlations account for hot-spot residues that are distributed in both N-terminal loops and protease core. Here we note that residues Ile19, Tyr20, and Leu24 were common to both SPM and node degeneracy calculations indicating these residues as critical during the allosteric regulation. Our results are in agreement with structural studies^100^, which have shown that large non-polar side chains of Ile are critical for the integrity of the N-terminal loops.

Our detailed analysis of intra- and inter-ring allosteric pathways reveals stronger communication in the open configuration between each ADEP binding site and N-terminal loops of distant protomers. According to these results, in this configuration neighboring intra-ring protomers are strongly coupled, consistent with the observed ability of the ATPase to trigger gate opening even as it activates only six binding sites. Interestingly, inter-ring coupling is also strengthened in the open configuration, even as the handle interface present in the closed configuration between protomer partners is removed. Stronger coupling in the open configuration is effected through efficient crossing of a single gap between residue communities that reduces the penalties of crossing gaps between multiple communities in the closed configuration.

Allosteric communication in the ClpP peptidase is likely to be further modulated by two external factors, namely its interactions with the ATPase partner, such as the single-ring ClpX or the double-ring ClpA, and with the substrate protein being degraded. The effect of the ATPase interaction reflects the variability of IGL/IGF loop binding to the seven ClpP binding sites during the catalytic cycle. Given the asymmetric binding of the six ATPase loops to the seven binding sites of ClpP, it is plausible that the signaling induced by ADEP binding represents the upper bound to the coupling strength between binding sites and the ClpP N-terminal loops. Additionally, the non-concerted conformational transitions of the ATPase hexamer further weakens the allosteric coupling and breaks the ring symmetry. Nevertheless, asymmetric intra-ring allostery may support ClpP’s active internalization of the polypeptide chain in the degradation process through non-concerted conformational changes of the pore loops. In support of the active action, studies using a ClpAP complex with one or more IGL loops of ClpA covalently crosslinked to the ClpP binding sites allow degradation to proceed at a slightly reduced rate compared with the noncovalent ClpAP complex^101^.

Allosteric communication within the ATPase itself has a high complexity and can therefore give rise to multiple responses in ClpP. An illustration of the complex ATPase allostery is provided by the double-ring ClpB disaggregase, which is not a cellular partner of ClpP, but can be engineered to form a complex with it^102^. Our community network analysis of apo, nucleotide and/or substrate-bound configurations of ClpB revealed distinct intra-ring communication within the nucleotide-binding domain (NBD) 1 and 2 rings^90^. Whereas, in the NBD1 ring, strong coupling is found between Large and Small subdomains of neighboring subunits, in the NBD2 ring, intra-protomer coupling is dominant.

The interaction between the ClpP peptidase and its substrate protein partner also has the potential to effect changes in the allosteric signaling. As noted in a recent study, ClpP forcefully grips the titin substrates with forces that exceed those of the partner ATPase^103^. Such strong signaling may dramatically alter the allosteric paths and break the symmetry of communications between the ADEP binding site and N-terminal loops.

## SUPPLEMENTARY MATERIAL

See the supplementary material for Table S1 showing the summary of the MD setups for ClpP wild-type and mutant setups, Table S2 for hot-spot residues derived from the structural perturbation method, Tables S3-S5 for highly correlated residue communities and Tables S6-S7 for the list of optimal paths in each ClpP configuration, Fig. S1 for RMSD time series in each ClpP configuration, Fig. S2 for DCCM convergence computed over multiple trajectories, Fig. S3 for normalized amplitudes of amino-acid motions associated with the top 5 normal modes, Fig. S4 for motions associated with top normal modes of the ClpP tetradecamer, Fig. S5 for the largest 20 eigenvalues of the PC modes for each ClpP configuration, Fig. S6 for network modularity, Fig. S7 for path length distributions of equidistant protomer loops, Movies SM1-4 (Multimedia view) for motions corresponding to principal components in the open (SM1–2) and closed pore configurations (SM3–4) and Movies SM5-6 (Multimedia view) for motions associated with the top two normal modes.

## ACKNOWLEDGMENTS

The authors gratefully acknowledge stimulating discussions with Sue Wickner and Mike Maurizi. This work has been supported by the National Science Foundation grants MCB-1516918 and MCB-2136816 to G.S. This work used the Extreme Science and Engineering Discovery Environment (XSEDE), which is supported by NSF grant number ACI-1548562. XSEDE Bridges resources at the Pittsburgh Supercomputer Center were used through allocation TG-MCB170020 to G.S. Partial support was received through research cyberinfrastructure resources and services provided by the Advanced Research Computing center at the University of Cincinnati.

## Data availability

The data that support the findings of this study are available from the corresponding author upon reasonable request.

## REFERENCES

1 S. Wickner, M. R. Maurizi, and S. Gottesman, Science 286, 1888 (1999).

2 M. Gersch, K. Famulla, M. Dahmen, C. Göbl, I. Malik, K. Richter, V. S. Korotkov, P. Sass, H. Rübsamen-Schaeff, and T. Madl, Nat. Commun. 6, 6320 (2015).

3 J. Ortega, H. S. Lee, M. R. Maurizi, and A. C. Steven, J. Struct. Biol. 146, 217 (2004).

4 J. Wang, J. A. Hartling, and J. M. Flanagan, Cell 91, 447 (1997).

5 X. Fei, T. A. Bell, S. Jenni, B. M. Stinson, T. A. Baker, S. C. Harrison, and R. T. Sauer, eLife 9, e52774 (2020).

6 Y. Katayama-Fujimura, S. Gottesman, and M. Maurizi, J. Biol. Chem. 262, 4477 (1987).

7 M. R. Maurizi, W. P. Clark, Y. Katayama, S. Rudikoff, J. Pumphrey, B. Bowers, and S. Gottesman, J. Biol. Chem. 265, 12536 (1990).

8 K. M. Woo, W. J. Chung, D. B. Ha, A. Goldberg, and C. H. Chung, J. Biol. Chem. 264, 2088 (1989).

9 M. W. Thompson, S. K. Singh, and M. R. Maurizi, J. Biol. Chem. 269, 18209 (1994).

10 S. Gottesman, M. R. Maurizi, and S. Wickner, Cell 91, 435 (1997).

11 K. R. Schmitz, D. W. Carney, J. K. Sello, and R. T. Sauer, Proc. Natl. Acad. Sci. U.S.A. 111, E4587 (2014).

12 M. E. Lee, T. A. Baker, and R. T. Sauer, J. Mol. Biol. 399, 707 (2010).

13 M. W. Thompson and M. R. Maurizi, J. Biol. Chem. 269, 18201 (1994).

14 J. A. Alexopoulos, A. Guarné, and J. Ortega, J. Struct. Biol. 179, 202 (2012).

15 A. Martin, T. A. Baker, and R. T. Sauer, Mol. Cell 27, 41 (2007).

16 G. Effantin, M. R. Maurizi, and A. C. Steven, J. Biol. Chem. 285, 14834 (2010).

17 A. Martin, T. A. Baker, and R. T. Sauer, Nat. Struct. Mol. Biol. 15, 1147 (2008).

18 C. Lee, M. P. Schwartz, S. Prakash, M. Iwakura, and A. Matouschek, Mol. cell 7, 627 (2001).

19 M.-E. Aubin-Tam, A. O. Olivares, R. T. Sauer, T. A. Baker, and M. J. Lang, Cell 145, 257 (2011).

20 R. A. Maillard, G. Chistol, M. Sen, M. Righini, J. Tan, C. M. Kaiser, C. Hodges, A. Martin, and C. Bustamante, Cell 145, 459 (2011).

21 A. Kravats, M. Jayasinghe, and G. Stan, Proc. Natl. Acad. Sci. U.S.A. 108, 2234 (2011).

22 A. O. Olivares, A. R. Nager, O. Iosefson, R. T. Sauer, and T. A. Baker, Nat. Struct. Mol. Biol. 21, 871 (2014).

23 A. Javidialesaadi, S. M. Flournoy, and G. Stan, J. Phys. Chem. B 123, 2623 (2019).

24 H. Y. Y. Fonseka, A. Javidi, L. F. Oliveira, C. Micheletti, and G. Stan, J. Phys. Chem. B 125, 7335 (2021).

25 R. A. Varikoti, H. Y. Y. Fonseka, M. S. Kelly, A. Javidi, M. Damre, S. Mullen, J. L. Nugent, C. M. Gonzales, G. Stan, and R. I. Dima, Nanomaterials 12, 1849 (2022).

26 L. D. Jennings, J. Bohon, M. R. Chance, and S. Licht, Biochemistry 47, 11031 (2008).

27 S. G. Kang, M. R. Maurizi, M. Thompson, T. Mueser, and B. Ahvazi, J. Struct. Biol. 148, 338 (2004).

28 A. Gribun, M. S. Kimber, R. Ching, R. Sprangers, K. M. Fiebig, and W. A. Houry, J. Biol. Chem. 280, 16185 (2005).

29 M. C. Bewley, V. Graziano, K. Griffin, and J. M. Flanagan, J. Struct. Biol. 153, 113 (2006).

30 Y.-I. Kim, I. Levchenko, K. Fraczkowska, R. V. Woodruff, R. T. Sauer, and T. A. Baker, Nat. Struct. Mol. Biol. 8, 230 (2001).

31 D. Y. Kim and K. K. Kim, J. Biol. Chem. 278, 50664 (2003).

32 Ž. Maglica, K. Kolygo, and E. Weber-Ban, Structure 17, 508 (2009).

33 A. J. Amor, K. R. Schmitz, J. K. Sello, T. A. Baker, and R. T. Sauer, ACS Chem. Biol. 11, 1552 (2016).

34 D. H. S. Li, Y. S. Chung, M. Gloyd, E. Joseph, R. Ghirlando, G. D. Wright, Y.-Q. Cheng, M. R. Maurizi, A. Guarne, and J. Ortega, Chem. Biol. 17, 959 (2010).

35 H. Brötz-Oesterhelt, D. Beyer, H.-P. Kroll, R. Endermann, C. Ladel, W. Schroeder, B. Hinzen, S. Raddatz, H. Paulsen, K. Henninger, J. E. Bandow, H.-G. Sahl, and H. Labischinski, Nat. Med. 11, 1082 (2005).

36 M. C. Bewley, V. Graziano, K. Griffin, and J. M. Flanagan, J. Struct. Biol. 165, 118 (2009).

37 S. Vahidi, Z. A. Ripstein, M. Bonomi, T. Yuwen, M. F. Mabanglo, J. B. Juravsky, K. Rizzolo, A. Velyvis, W. A. Houry, M. Vendruscolo, J. L. Rubinstein, and L. E. Kay, Proc. Natl. Acad. Sci. U.S.A. 115, E6447 (2018).

38 J. Kirstein, A. Hoffmann, H. Lilie, R. Schmidt, H. Rübsamen-Waigmann, H. Brötz-Oesterhelt, A. Mogk, and K. Turgay, EMBO Mol. Med. 1, 37 (2009).

39 P. Sass, M. Josten, K. Famulla, G. Schiffer, H.-G. Sahl, L. Hamoen, and H. Brötz-Oesterhelt, Proc. Natl. Acad. Sci. U.S.A. 108, 17474 (2011).

40 R. M. Raju, A. L. Goldberg, and E. J. Rubin, Nat. Rev. Drug Discov. 11, 777 (2012).

41 B. P. Conlon, E. S. Nakayasu, L. E. Fleck, M. D. LaFleur, V. M. Isabella, K. Coleman, S. N. Leonard, R. D. Smith, J. N. Adkins, and K. Lewis, Nature 503, 365 (2013).

42 J. Zhang, F. Ye, L. Lan, H. Jiang, C. Luo, and C.-G. Yang, J. Biol. Chem. 286, 37590 (2011).

43 O. Keskin, I. Bahar, D. Flatow, D. G. Covell, and R. L. Jernigan, Biochemistry 41, 491 (2002).

44 W. Zheng, B. R. Brooks, and D. Thirumalai, Biophys. J. 93, 2289 (2007).

45 M. Jayasinghe, P. Shrestha, X. Wu, R. Tehver, and G. Stan, Biophys. J. 103, 1285 (2012).

46 R. Gruber and A. Horovitz, Chem. Rev. 116, 6588 (2016).

47 R. Gruber, M. Levitt, and A. Horovitz, Proc. Natl. Acad. Sci. U.S.A. 114, 5189 (2017).

48 A. R. Atilgan and C. Atilgan, Brief. Funct. Genomics 11, 479 (2012).

49 R. Nussinov, C. J. Tsai, and B. Ma, Annu. Rev. Biophys. 42, 169 (2013).

50 V. A. Feher, J. D. Durrant, A. T. Van Wart, and R. E. Amaro, Curr. Opin. Struct. Biol. 25, 98 (2014).

51 J. Guo and H.-X. Zhou, Chem. Rev. 116, 6503 (2016).

52 A. R. Atilgan, P. Akan, and C. Baysal, Biophys. J. 86, 85 (2004).

53 C. Chennubhotla and I. Bahar, Mol. Syst. Biol. 2, e172 (2006).

54 A. tSethi, J. Eargle, A. A. Black, and Z. Luthey-Schulten, Proc. Natl. Acad. Sci. U.S.A. 106, 6620 (2009).

55 I. Rivalta, M. M. Sultan, N. S. Lee, G. A. Manley, J. P. Loria, and V. S. Batista, Proc. Natl. Acad. Sci. U.S.A. 109, E1428 (2012).

56 Y. Miao, S. E. Nichols, P. M. Gasper, V. T. Metzger, and J. A. McCammon, Proc. Natl. Acad. Sci. U.S.A. 110, 10982 (2013).

57 P. M. Gasper, B. Fuglestad, E. A. Komives, P. R. Markwick, and J. A. McCammon, Proc. Natl. Acad. Sci. U.S.A. 109, 21216 (2012).

58 G. Scarabelli and B. J. Grant, Biophys. J. 107, 2204 (2014).

59 A. T. Van Wart, J. Durrant, L. Votapka, and R. E. Amaro, J. Chem. Theory Comput. 10, 511 (2014).

60 G. Palermo, C. G. Ricci, A. Fernando, R. Basak, M. Jinek, I. Rivalta, V. S. Batista, and J. A. McCammon, J. Am. Chem. Soc. 139, 16028 (2017).

61 G. Stetz and G. M. Verkhivker, PLoS Comput. Biol. 13, e1005299 (2017).

62 G. M. Verkhivker and L. D. Paola, J. Phys. Chem. B 125, 850 (2021).

63 W. Humphrey, A. Dalke, and K. Schulten, J. Mol. Graph. 14, 33 (1996).

64 A. Fiser and A. Sali, Bioinformatics 19, 2500 (2003).

65 Schrödinger, LLC, “The PyMOL molecular graphics system, version 1.8,” (2015).

66 D. Van Der Spoel, E. Lindahl, B. Hess, G. Groenhof, A. E. Mark, and H. J. Berendsen, J. Comput. Chem. 26, 1701 (2005).

67 R. B. Best, X. Zhu, J. Shim, P. E. Lopes, J. Mittal, M. Feig, and A. D. MacKerell Jr, J. Chem. Theory Comput. 8, 3257 (2012).

68 K. Vanommeslaeghe, E. Hatcher, C. Acharya, S. Kundu, S. Zhong, J. Shim, E. Darian, O. Guvench, P. Lopes, I. Vorobyov, et al., J. Comput. Chem. 31, 671 (2010).

69 E. Neria, S. Fischer, and M. Karplus, J. Chem. Phys. 105, 1902 (1996).

70 G. Bussi, D. Donadio, and M. Parrinello, J. Chem. Phys. 126, 014101 (2007).

71 M. Parrinello and A. Rahman, J. Appl. Phys. 52, 7182 (1981).

72 B. J. Grant, A. P. Rodrigues, K. M. ElSawy, J. A. McCammon, and L. S. Caves, Bioinformatics 22, 2695 (2006).

73 H. Kamberaj and A. Van der Vaart, Biophys. J. 96, 1307 (2009).

74 M. Girvan and M. E. J. Newman, Proc. Natl. Acad. Sci. U.S.A. 99, 7821 (2002).

75 B. J. Grant, L. Skjærven, and X.-Q. Yao, Protein Science 30, 20 (2021).

76 S. Hayward and B. L. De Groot, in Molecular Modeling of Proteins (Springer, 2008) pp. 89–106.

77 A. Amadei, A. B. Linssen, and H. J. Berendsen, Proteins 17, 412 (1993).

78 M. A. Balsera, W. Wriggers, Y. Oono, and K. Schulten, J. Phys. Chem. 100, 2567 (1996).

79 A. Amadei, B. De Groot, M.-A. Ceruso, M. Paci, A. Di Nola, and H. Berendsen, Proteins 35, 283 (1999).

80 R. Cossio-Pérez, J. Palma, and G. Pierdominici-Sottile, J. Chem. Inf. Model 57, 826 (2017).

81 X.-Q. Yao, R. U. Malik, N. W. Griggs, L. Skjærven, J. R. Traynor, S. Sivaramakrishnan, and B. J. Grant, J. Biol. Chem. 291, 4742 (2016).

82 L. E. Bradley, “‘Overlapping Coefficient’. In Encyclopedia of Statistical Sciences.” (John Wiley: New York, 1985) Chap. 6, pp. 546–547.

83 M. S. Weitzman, “Measures of overlap of income distributions of white and negro families in the United States,” (Vol. 22. US Bureau of the Census, 1970).

84 M. M. Tirion, Phys. Rev. Lett. 77, 1905 (1996).

85 I. Bahar and A. Rader, Curr. Opin. Struct. Biol. 15, 586 (2005).

86 W. Zheng, B. R. Brooks, and D. Thirumalai, Proc. Natl. Acad. Sci. U.S.A. 103, 7664 (2006).

87 R. Tehver, J. Chen, and D. Thirumalai, J. Mol. Biol. 387, 390 (2009).

88 O. Marques and Y.-H. Sanejouand, Proteins 23, 557 (1995).

89 L. Skjaerven, A. Martinez, and N. Reuter, Proteins 79, 232 (2011).

90 M. Damre, A. Dayananda, R. A. Varikoti, G. Stan, and R. I. Dima, Biophys. J. 120, 3437 (2021).

91 M. T. Posgai, S. Tonddast-Navaei, M. Jayasinghe, G. M. Ibrahim, G. Stan, and A. B. Herr, Proc. Natl. Acad. Sci. U.S.A. 115, E8882 (2018).

92 L. González-Paz, C. Lossada, M. L. Hurtado-León, F. V. Fernández-Materán, J. L. Paz, S. Parvizi, R. E. Cardenas Castillo, F. Romero, and Y. J. Alvarado, ACS Omega, in press (2023).

93 O. F. Lange and H. Grubmüller, Proteins: Struct., Funct., and Bioinf. 62, 1053 (2006).

94 L. G. Ahuja, S. S. Taylor, and A. P. Kornev, IUBMB life 71, 685 (2019).

95 M. C. Melo, R. C. Bernardi, C. De La Fuente-Nunez, and Z. Luthey-Schulten, J. Chem. Phys. 153, 134104 (2020).

96 L. Astl, G. Stetz, and G. M. Verkhivker, J. Chem. Inf. Model. 60, 3616 (2020).

97 Y. Yuan, J. Deng, and Q. Cui, J. Am. Chem. Soc. 144, 10870 (2022).

98 M. Iljina, H. Mazal, A. Dayananda, Z. Zhang, G. Stan, I. Riven, and G. Haran, bioRxiv (2023), 10.1101/2023.02.02.526786.

99 V. Bhandari, K. S. Wong, J. L. Zhou, M. F. Mabanglo, R. A. Batey, and W. A. Houry, ACS Chem. Biol. 13, 1413 (2018).

100 J. Alexopoulos, B. Ahsan, L. Homchaudhuri, N. Husain, Y.-Q. Cheng, and J. Ortega, Mol. Microbiol. 90, 167 (2013).

101 S. Kim, K. L. Zuromski, T. A. Bell, R. T. Sauer, and T. A. Baker, eLife 9, e61451 (2020).

102 J. Weibezahn, C. Schlieker, B. Bukau, and A. Mogk, J. Biol. Chem. 278, 32608 (2003).

103 S. D. Walker and A. O. Olivares, Biophys. J. 121, 3907 (2022).

