## Supplemental Tables and Figures for "Allosteric Communication in the Gating Mechanism for Controlled Protein Degradation by the Bacterial ClpP Peptidase"

*Affiliations:*

### Supplementary Tables

**Table S1.** Summary of the MD setups for ClpP wt and mutant setups

| State | Setup | # Atoms | # Waters | # Na Ions |
|---|---|---|---|---|
| 3MT6 | WT | 222757 | 59585 | 70 |
| | WT(ΔADEP ) | 222496 | 60002 | 70 |
| | E8K | 222605 | 60015 | 42 |
| | E14A-R15A | 222404 | 60060 | 70 |
| | E14A-R15A-E8A | 222434 | 60098 | 56 |
| | E14A-R15A-K25A | 222343 | 60091 | 84 |
| | I7P | 222435 | 60005 | 70 |
| | K25E | 222387 | 59989 | 98 |
| 1YG6 | WT | 209623 | 55711 | 70 |
| | E8K | 209693 | 55711 | 42 |
| | E14A-R15A | 209546 | 55774 | 70 |
| | E14A-R15A-E8A | 209498 | 55786 | 56 |
| | E14A-R15A-K25A | 209422 | 55784 | 84 |
| | I7P | 209589 | 55723 | 70 |
| | K25E | 209502 | 55694 | 98 |

**Table S2.** Hot-spot residues derived from the structural perturbation values for the ClpP. Hot-spot residues were obtained for the top two overlap modes. Bold are the common residues for the two modes

| Mode 38 | Mode 28 |
|---|---|
| **VAL3** | **VAL3** |
| PRO4 | **MET5** |
| **MET5** | **VAL6** |
| **VAL6** | **ILE7** |
| **ILE7** | **GLU8** |
| **GLU8** | **GLN9** |
| **GLN9** | **THR10** |
| **THR10** | SER11 |
| ARG12 | **GLY13** |
| **GLY13** | **GLU14** |
| **GLU14** | **ARG15** |
| **ARG15** | **SER16** |
| **SER16** | **PHE17** |
| **PHE17** | **ASP18** |
| **ASP18** | ILE19 |
| **TYR20** | **TYR20** |
| **ARG22** | **ARG22** |
| LEU23 | **VAL28** |
| LEU24 | HIS191 |
| LYS25 | ARG192 |
| **VAL28** | |

**Table S3.** Highly correlated residue communities in the ClpP open configuration

| Community ID | Number of residues | Residue segments |
|---|---|---|
| 1 | 193 | c(1:124, 129:193, 1476:1479) |
| 2 | 47 | c(125:128, 1480:1522) |
| 3 | 193 | c(194:270, 272:316, 321:386, 1667:1671) |
| 4 | 148 | c(271, 387:508, 542, 557:576, 693:696) |
| 5 | 189 | c(317:320, 1545:1666, 1672:1734) |
| 6 | 95 | c(509:541, 543:556, 1860:1907) |
| 7 | 3 | 577:579 |
| 8 | 144 | c(580:652, 654:692, 697:701, 734:736, 749:772) |
| 9 | 191 | c(653, 773:894, 901:962, 2245:2250) |
| 10 | 93 | c(702:733, 737:748, 2052:2100) |
| 11 | 193 | c(895:900, 2124:2244, 2251:2316) |
| 12 | 3 | 963:965 |
| 13 | 149 | c(966:1088, 1121, 1135:1158, 1274) |
| 14 | 58 | c(1089:1097, 2439:2487) |
| 15 | 36 | c(1098:1120, 1122:1134) |
| 16 | 143 | c(1159:1273, 1275:1279, 1329:1351) |

| | | |
|---|---|---|
| 17 | 183 | c(1280:1290, 1293, 2532:2702) |
| 18 | 37 | c(1291:1292, 1294:1328) |
| 19 | 164 | c(1352:1475, 1523:1541, 2511:2531) |
| 20 | 3 | 1542:1544 |
| 21 | 3 | 1735:1737 |
| 22 | 145 | c(1738:1859, 1908:1930) |
| 23 | 1 | 1931 |
| 24 | 140 | c(1932:2051, 2101:2120) |
| 25 | 3 | 2121:2123 |
| 26 | 144 | c(2317:2438, 2488:2509) |
| 27 | 1 | 2510 |

**Table S4**. Highly correlated residue communities in the ClpP closed configuration

| Community ID | Number of residues | Residue segments |
|---|---|---|
| 1 | 144 | c(1:122, 172:193) |
| 2 | 95 | c(123:171, 1475:1520) |
| 3 | 1 | 194 |
| 4 | 21 | 195:215 |
| 5 | 121 | c(216:315, 363:383) |
| 6 | 95 | c(316:362, 1667:1714) |
| 7 | 3 | 384:386 |
| 8 | 27 | 387:413 |
| 9 | 116 | c(414:507, 558:579) |
| 10 | 98 | c(508:557, 1859:1906) |
| 11 | 142 | c(580:700, 749:769) |
| 12 | 64 | c(701:748, 2053:2067, 2070) |
| 13 | 3 | 770:772 |
| 14 | 142 | c(773:838, 840:849, 851:893, 943:965) |
| 15 | 1 | 839 |
| 16 | 117 | c(850, 992:1031, 1033:1086, 1136:1147, 1149:1158) |
| 17 | 96 | c(894:942, 2246:2292) |
| 18 | 30 | c(966:991, 1159:1162) |
| 19 | 1 | 1032 |
| 20 | 51 | c(1087:1093, 2444:2487) |
| 21 | 47 | c(1094:1135, 1148, 2440:2443) |

| | | |
|---|---|---|
| 22 | 141 | c(1163:1279, 1328:1351) |
| 23 | 93 | c(1280:1327, 2633:2664, 2666:2678) |
| 24 | 147 | c(1352:1428, 1430:1474, 1521:1544, 1622) |
| 25 | 146 | c(1429, 2510:2632, 2665, 2679:2699) |
| 26 | 1 | 1545 |
| 27 | 157 | c(1546:1621, 1623:1666, 1715:1737, 1806:1819) |
| 28 | 130 | c(1738:1803, 1805, 1820:1858, 1907:1930) |
| 29 | 1 | 1804 |
| 30 | 144 | c(1931:2052, 2102:2123) |
| 31 | 33 | c(2068:2069, 2071:2101) |
| 32 | 22 | 2124:2145 |
| 33 | 120 | c(2146:2189, 2191:2245, 2293:2313) |
| 34 | 1 | 2190 |
| 35 | 3 | 2314:2316 |
| 36 | 145 | c(2317:2439, 2488:2509) |
| 37 | 3 | 2700:2702 |

**Table S5.** Highly correlated residue communities in the ClpP open, no ADEP configuration

| Community ID | Number of residues | Residue segments |
|---|---|---|
| 1 | 152 | c(1:4, 1043, 1159:1280, 1315, 1328:1351) |
| 2 | 142 | c(5:66, 68:121, 169:182, 184:193, 309, 384) |
| 3 | 1 | 67 |
| 4 | 51 | c(122:128, 1478:1521) |
| 5 | 43 | c(129:168, 183, 1476:1477) |
| 6 | 161 | c(194:308, 310:315, 363:383, 385:402, 502) |
| 7 | 94 | c(316:362, 1667:1713) |
| 8 | 127 | c(403:451, 454:501, 503:508, 556:579) |
| 9 | 1 | 452 |
| 10 | 1 | 453 |
| 11 | 97 | c(509:555, 1859:1908) |
| 12 | 2 | 580:581 |
| 13 | 149 | c(582:701, 749:777) |
| 14 | 50 | c(702:708, 2057:2085, 2087:2100) |
| 15 | 44 | c(709:748, 2053:2056) |
| 16 | 139 | c(778:838, 840:894, 943:965) |
| 17 | 1 | 839 |
| 18 | 96 | c(895:942, 2245:2292) |
| 19 | 1 | 966 |
| 20 | 19 | c(967:984, 987) |

| | | |
|---|---|---|
| 21 | 122 | c(985:986, 988:1042, 1044:1087, 1135:1155) |
| 22 | 46 | c(1088:1091, 2444:2485) |
| 23 | 48 | c(1092:1134, 2439:2443) |
| 24 | 3 | 1156:1158 |
| 25 | 94 | c(1281:1314, 1316:1327, 2631:2665, 2667:2679) |
| 26 | 147 | c(1352:1475, 1522:1544) |
| 27 | 151 | c(1545:1548, 1550:1610, 1612:1666, 1714:1737, 1812, 1814:1819) |
| 28 | 136 | c(1549, 1739:1803, 1805:1811, 1813, 1820:1858, 1909:1930, 2008) |
| 29 | 1 | 1611 |
| 30 | 1 | 1738 |
| 31 | 1 | 1804 |
| 32 | 143 | c(1931:1934, 1936:1996, 1998:2007, 2009:2052, 2086, 2101:2123) |
| 33 | 168 | c(1935, 2124:2145, 2148, 2318:2431, 2433:2438, 2486:2509) |
| 34 | 1 | 1997 |
| 35 | 122 | c(2146:2147, 2149:2244, 2293:2316) |
| 36 | 1 | 2317 |
| 37 | 145 | c(2432, 2511:2630, 2666, 2680:2702) |
| 38 | 1 | 2510 |

**Table S6**. The optimal path between an individual ADEP binding site and the N-terminal loops in the cis ring in each ClpP configuration

| Configuration | Shortest Path |
|---|---|
| Open | Arg22 → Ile19 → Asp18 → Phe17 → Ser16 → Arg15 → Glu14 → Gly13 |
| Open, no ADEP | Arg22 → Phe17 → Ser16 → Glu14 → Gly13 |
| Closed | Arg22 → Phe17 → Ser16 → Arg15 → Glu14 → Gly13 |

**Table S7**. The optimal path between the ADEP binding site in protomer B of the cis ring and the N-terminal loop of its partner protomer I in the trans ring in each ClpP configuration.

| Configuration | Shortest Path |
|---|---|
| Open | B_Ile90 → B_Thr104 → B_Met153 → B_Asn150 → B_Lys146 → B_Leu143 → B_Ala139 → B_Ile135 → I_Leu125 → I_Pro124 → I_Gln123 → I_His122 → I_Ile121 → I_Val119 → I_Cys91 → I_Tyr62 → I_Val28 → I_Ser21 → I_Phe17 → I_Ser16 → I_Arg15 → I_Glu14 → I_Gly13 |
| Closed | B_Ile90 → B_Thr104 → B_His156 → B_Met153 → B_Asn150 → B_Lys146 → B_Leu143 → B_Ala139 → B_Ile135 → B_Gln131 → I_Gln123 → I_His122 → I_Met120 → I_Ala96 → I_Asn64 → I_Phe30 → I_Leu23 → I_Arg22 → I_Phe17 → I_Arg15 → I_Glu14 → I_Gly13 |

### Supplementary Figures

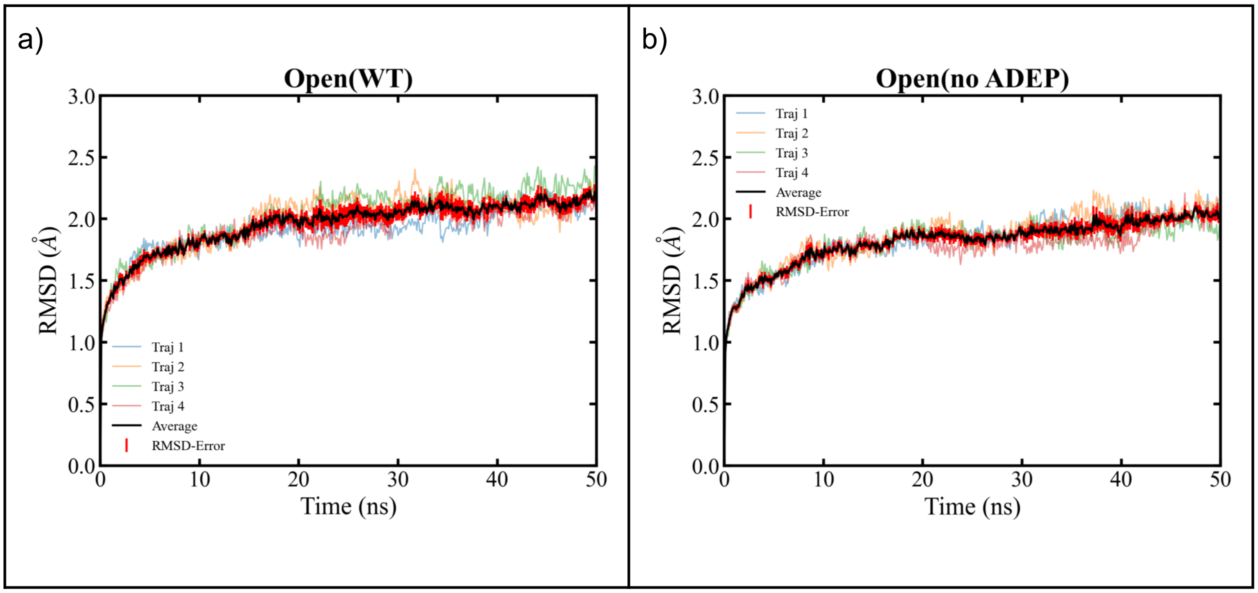

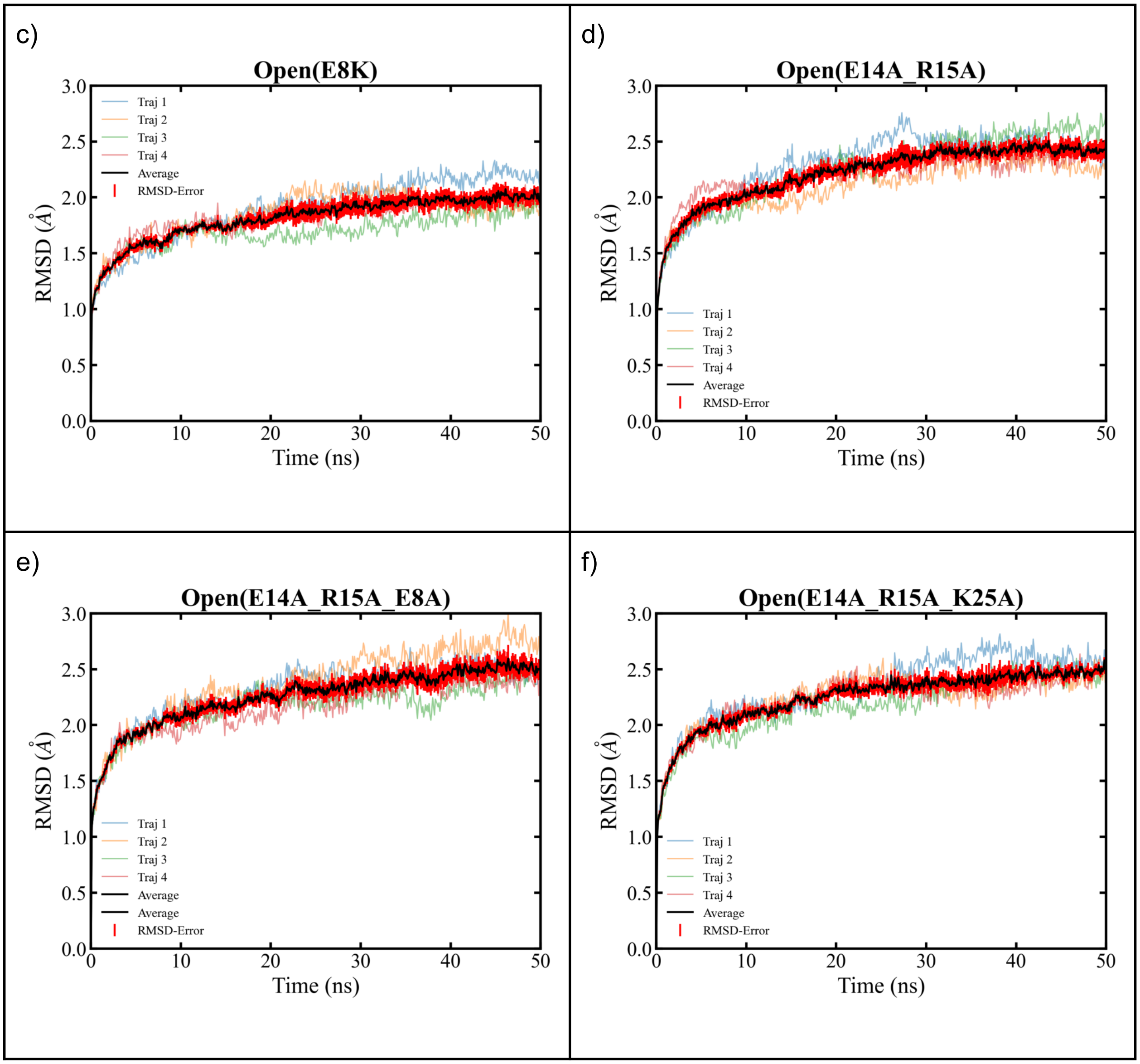

c)
Open(E8K)
RMSD (Å)
Time (ns)
Traj 1
Traj 2
Traj 3
Traj 4
Average
RMSD-Error
d)
Open(E14A_R15A)
RMSD (Å)
Time (ns)
Traj 1
Traj 2
Traj 3
Traj 4
Average
RMSD-Error
e)
Open(E14A_R15A_E8A)
RMSD (Å)
Time (ns)
Traj 1
Traj 2
Traj 3
Traj 4
Average
Average
RMSD-Error
f)
Open(E14A_R15A_K25A)
RMSD (Å)
Time (ns)
Traj 1
Traj 2
Traj 3
Traj 4
Average
RMSD-Error

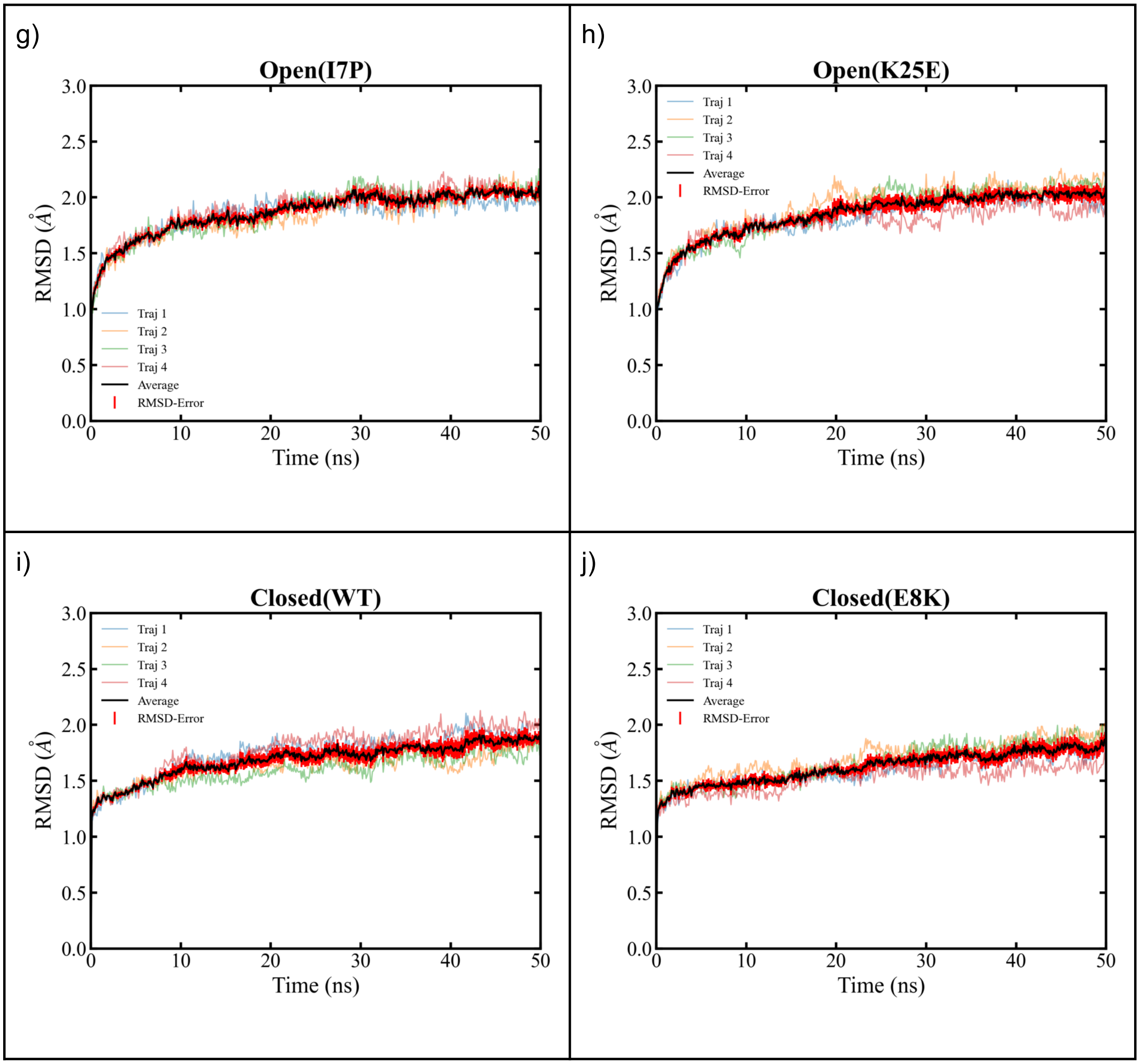
g)
Open(I7P)
h)
Open(K25E)
i)
Closed(WT)
j)
Closed(E8K)
RMSD (Å)
Time (ns)
Traj 1
Traj 2
Traj 3
Traj 4
Average
RMSD-Error
0.0
0.5
1.0
1.5
2.0
2.5
3.0
0
10
20
30
40
50

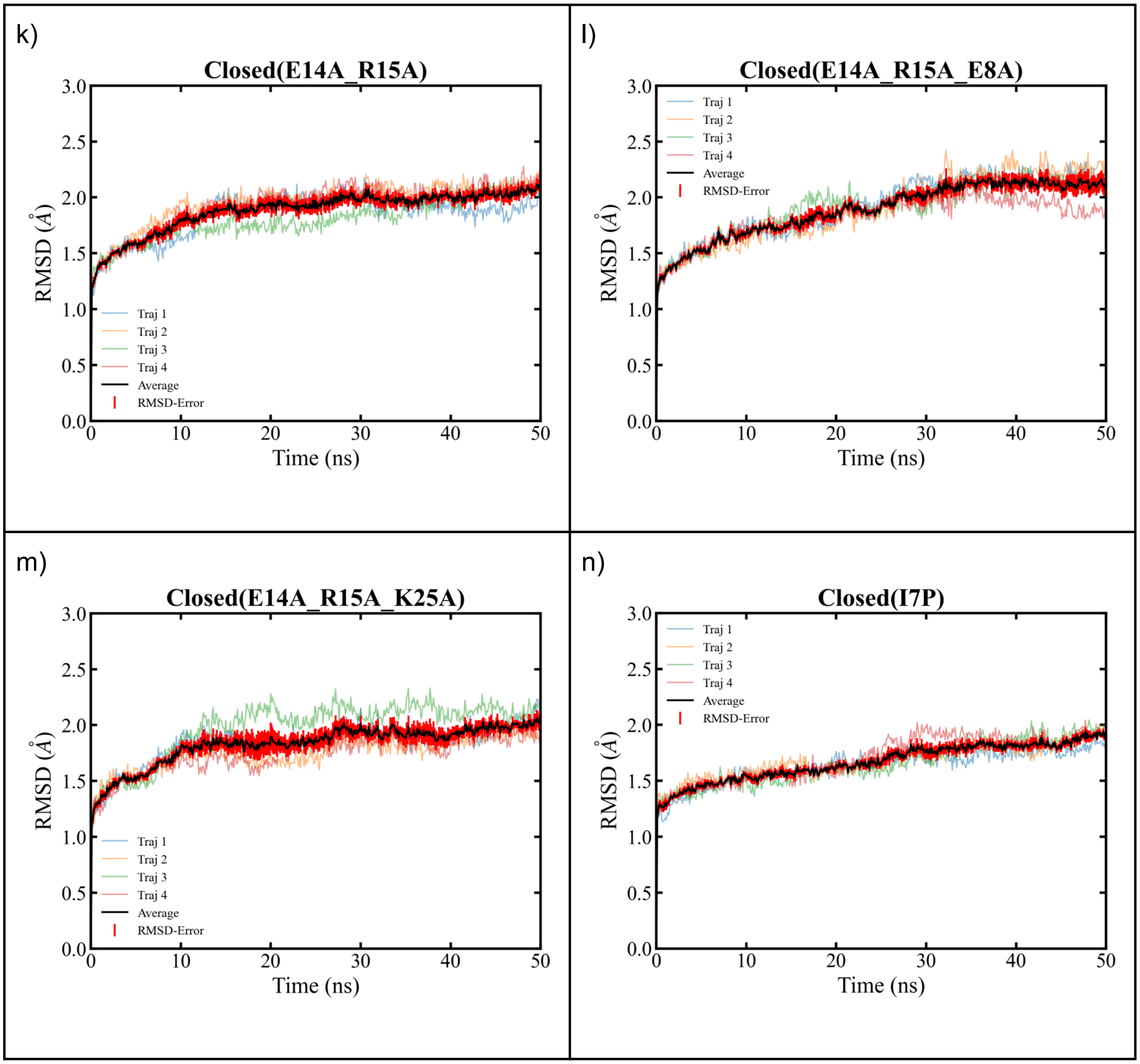
k)
Closed(E14A_R15A)
l)
Closed(E14A_R15A_E8A)
m)
Closed(E14A_R15A_K25A)
n)
Closed(I7P)
RMSD (Å)
Time (ns)
Traj 1
Traj 2
Traj 3
Traj 4
Average
RMSD-Error
0.0
0.5
1.0
1.5
2.0
2.5
3.0
0
10
20
30
40
50

o)

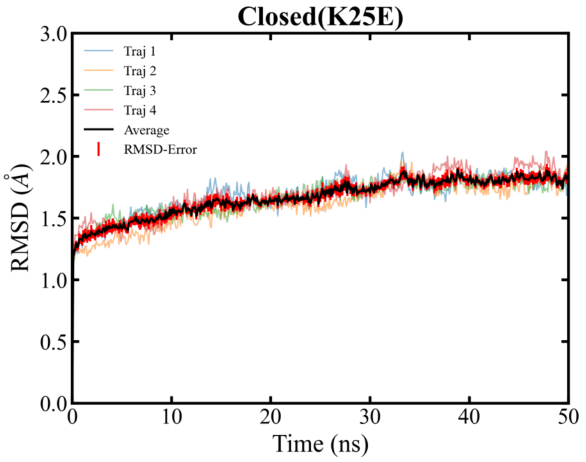


**Figure S1:** RMSD plots versus time, including ensemble averages (thick black lines) and standard errors (light red) over multiple trajectories for the various wild-type and mutant ClpP setups in (a)-(h) open, (i)-(o) closed pore configurations.

a)

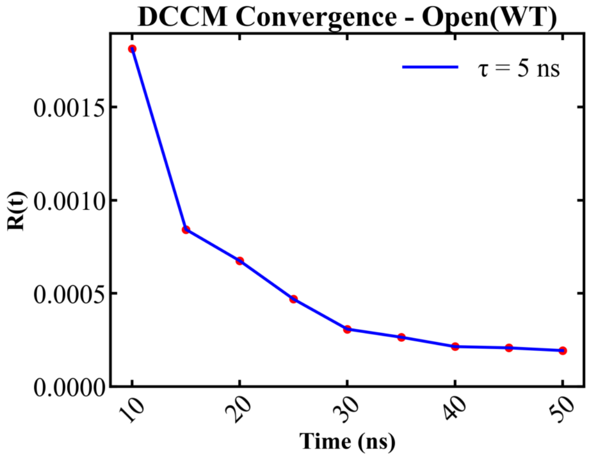


b)

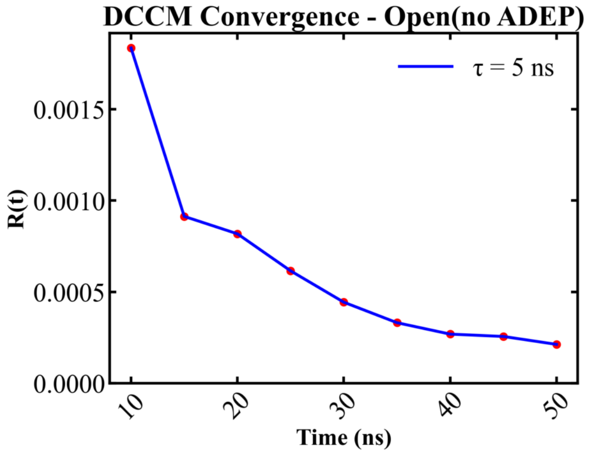

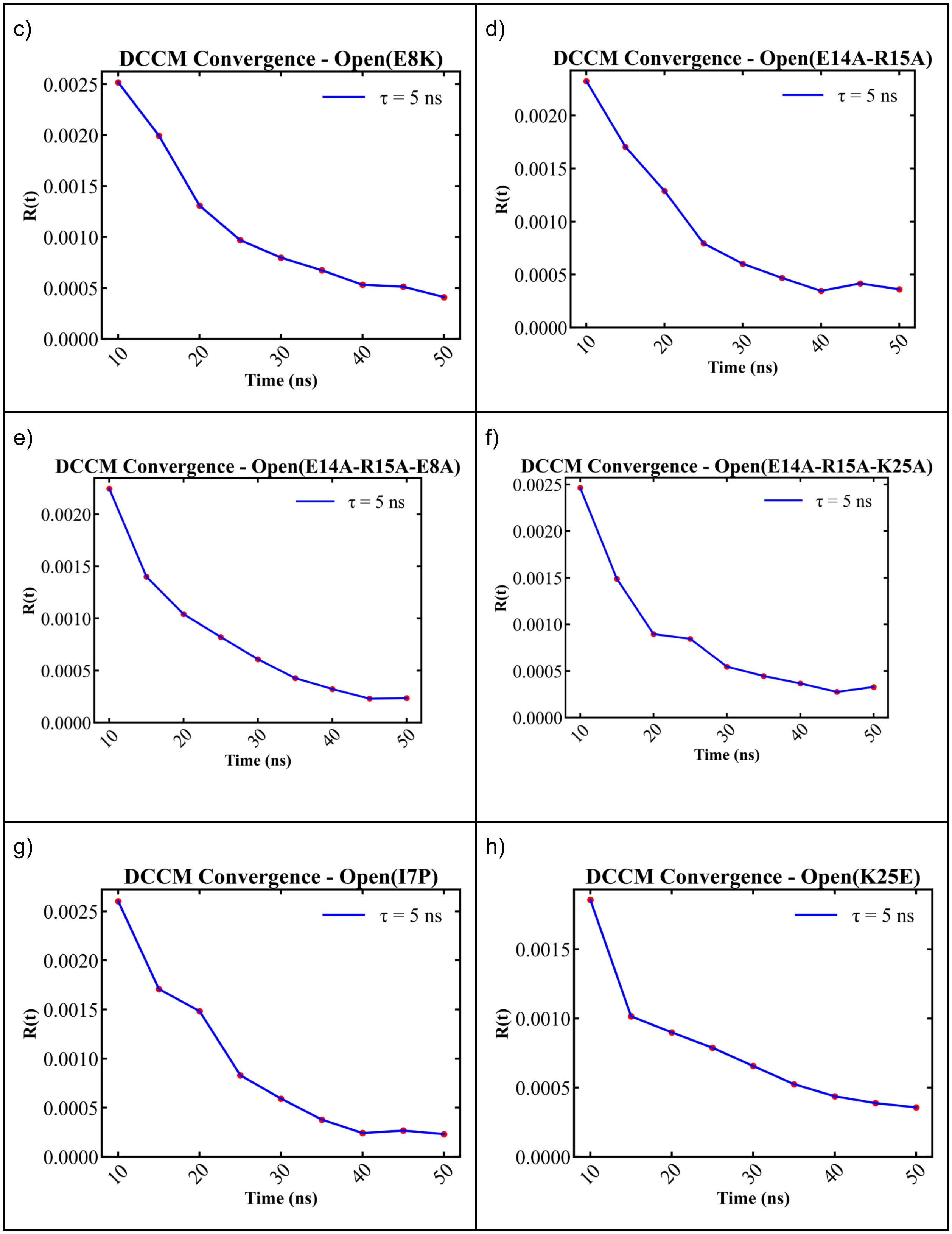

c)
DCCM Convergence - Open(E8K)
τ = 5 ns
R(t)
Time (ns)
d)
DCCM Convergence - Open(E14A-R15A)
τ = 5 ns
R(t)
Time (ns)
e)
DCCM Convergence - Open(E14A-R15A-E8A)
τ = 5 ns
R(t)
Time (ns)
f)
DCCM Convergence - Open(E14A-R15A-K25A)
τ = 5 ns
R(t)
Time (ns)
g)
DCCM Convergence - Open(I7P)
τ = 5 ns
R(t)
Time (ns)
h)
DCCM Convergence - Open(K25E)
τ = 5 ns
R(t)
Time (ns)

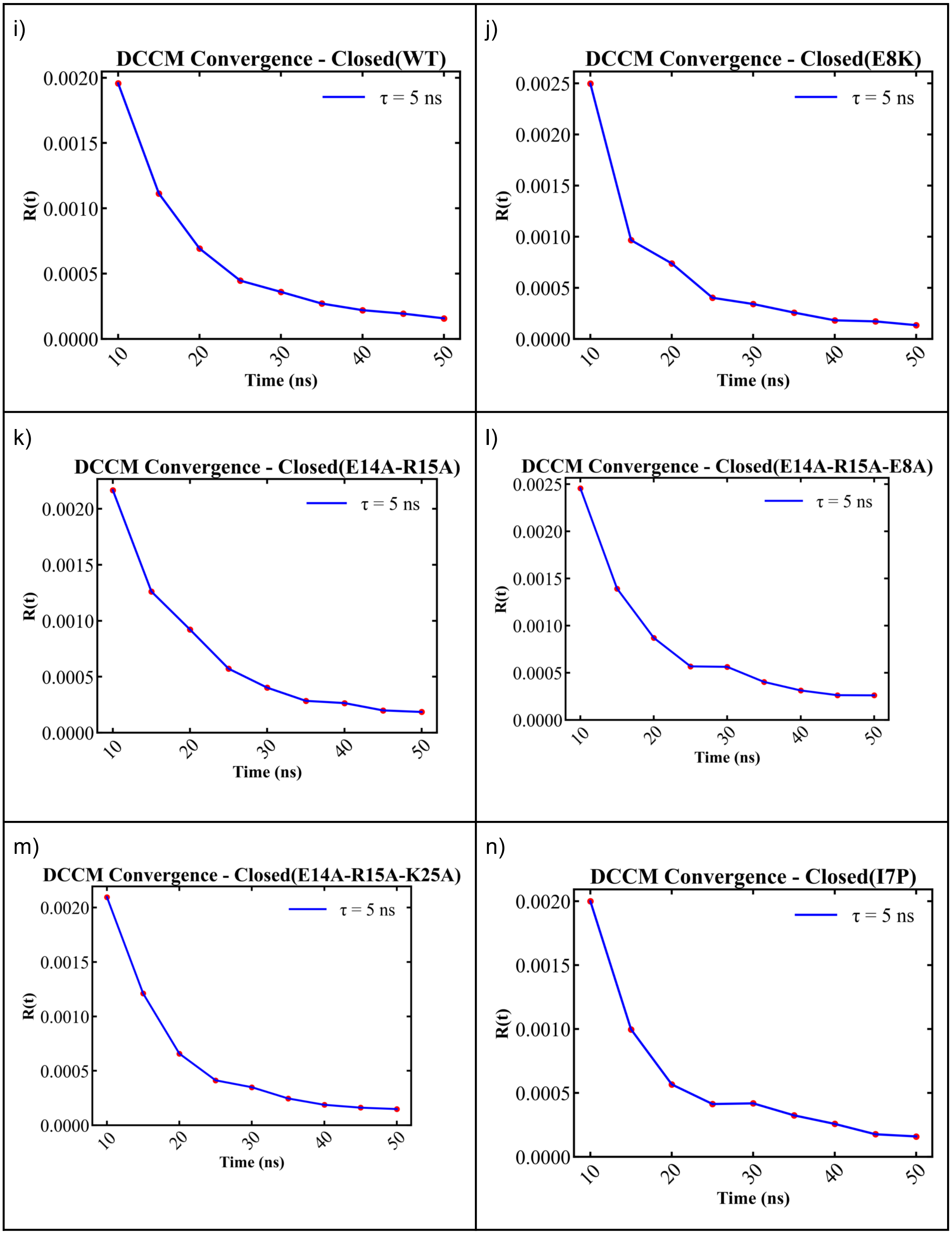
i)
DCCM Convergence - Closed(WT)
τ = 5 ns
R(t)
Time (ns)
j)
DCCM Convergence - Closed(E8K)
τ = 5 ns
R(t)
Time (ns)
k)
DCCM Convergence - Closed(E14A-R15A)
τ = 5 ns
R(t)
Time (ns)
l)
DCCM Convergence - Closed(E14A-R15A-E8A)
τ = 5 ns
R(t)
Time (ns)
m)
DCCM Convergence - Closed(E14A-R15A-K25A)
τ = 5 ns
R(t)
Time (ns)
n)
DCCM Convergence - Closed(I7P)
τ = 5 ns
R(t)
Time (ns)

o)

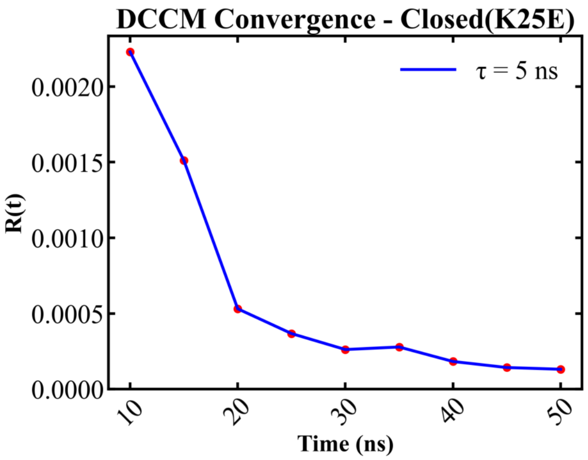


**Figure S2:** DCCM convergence computed over multiple trajectories for wild-type and mutant ClpP in (a)-(h) open (i)-(o) closed pore configurations.

A)

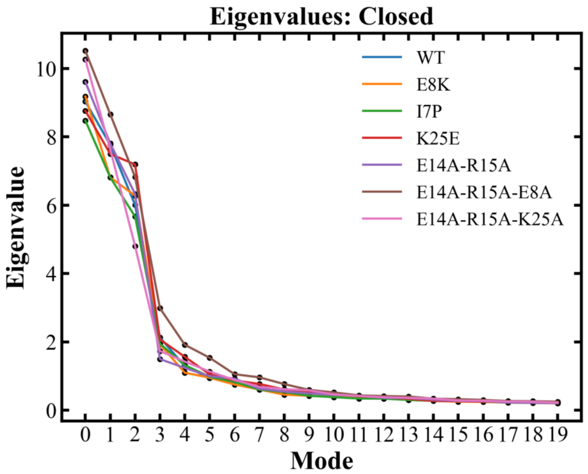


B)

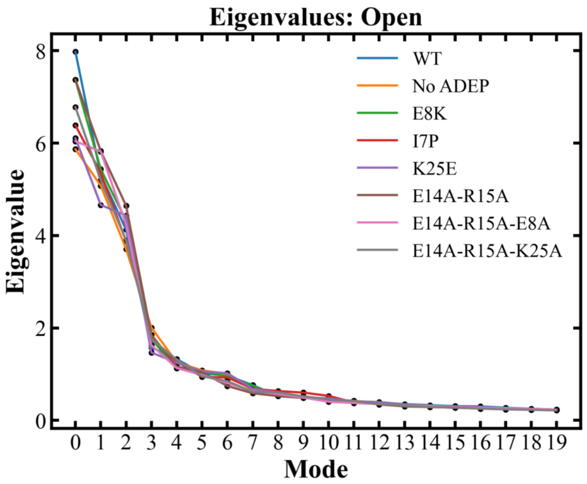


**Figure S3:** The largest 20 eigenvalues of the PC modes. A) Closed pore, wild-type and mutant ClpP B) Open, wild-type and mutant ClpP

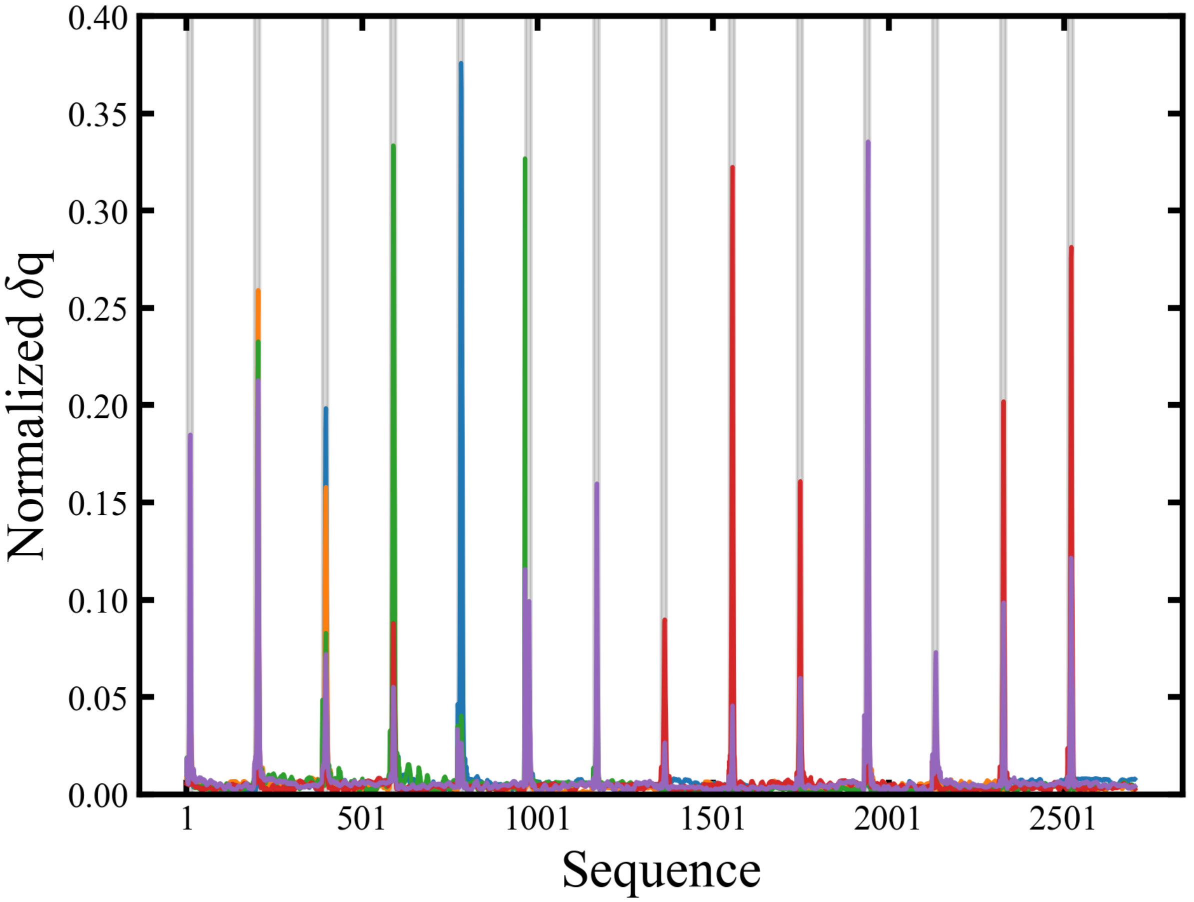


**Figure S4:** Normalized amplitudes of amino-acid motions associated with the top 5 normal modes (38, 27, 26, 25, 28) overlapped with the open → closed transition of ClpP. Shaded regions (gray) correspond to the N-terminal loop region (residues 1-18)

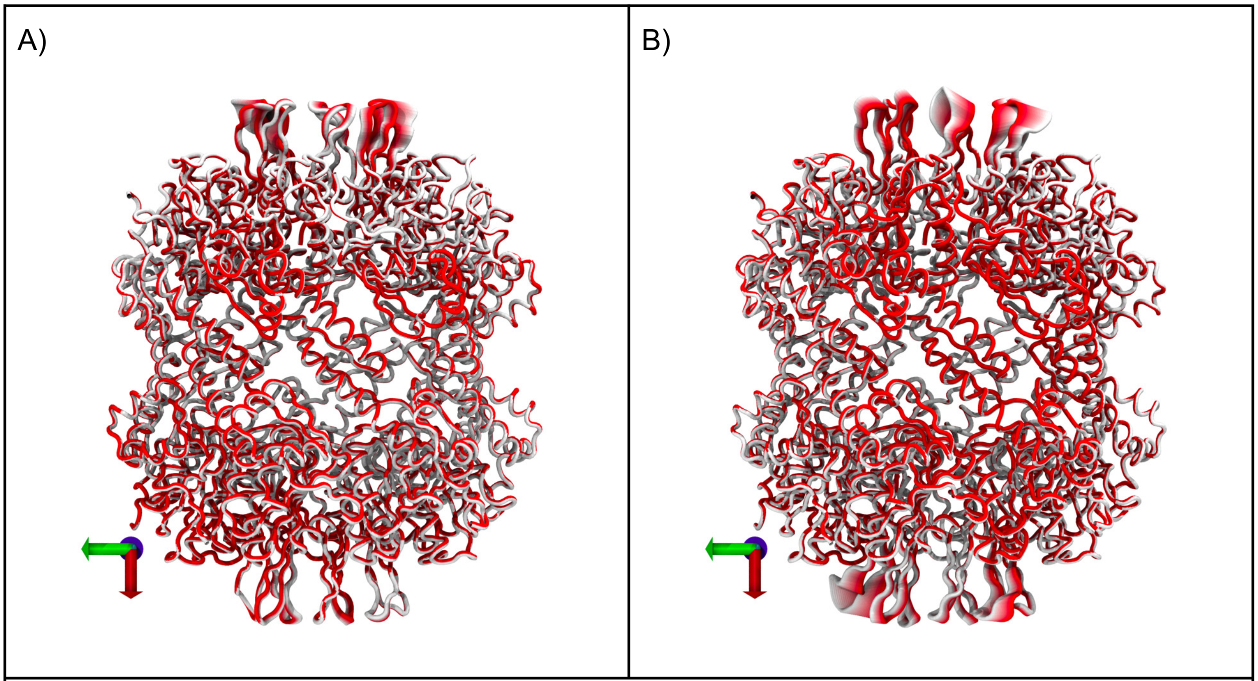


**Figure S5:** Motions associated with significant normal modes of the ClpP tetradecamer. A) Mode 38 B) Mode 28. The color gradient indicates locations with significant conformational changes.

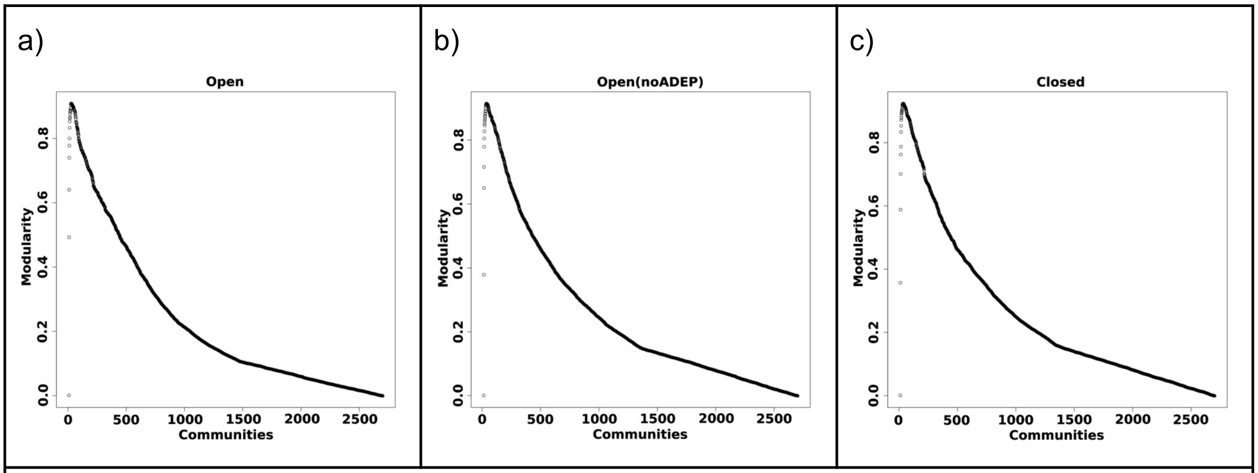


**Figure S6:** Network modularity in a) open b) open, no ADEP, c) closed ClpP configurations

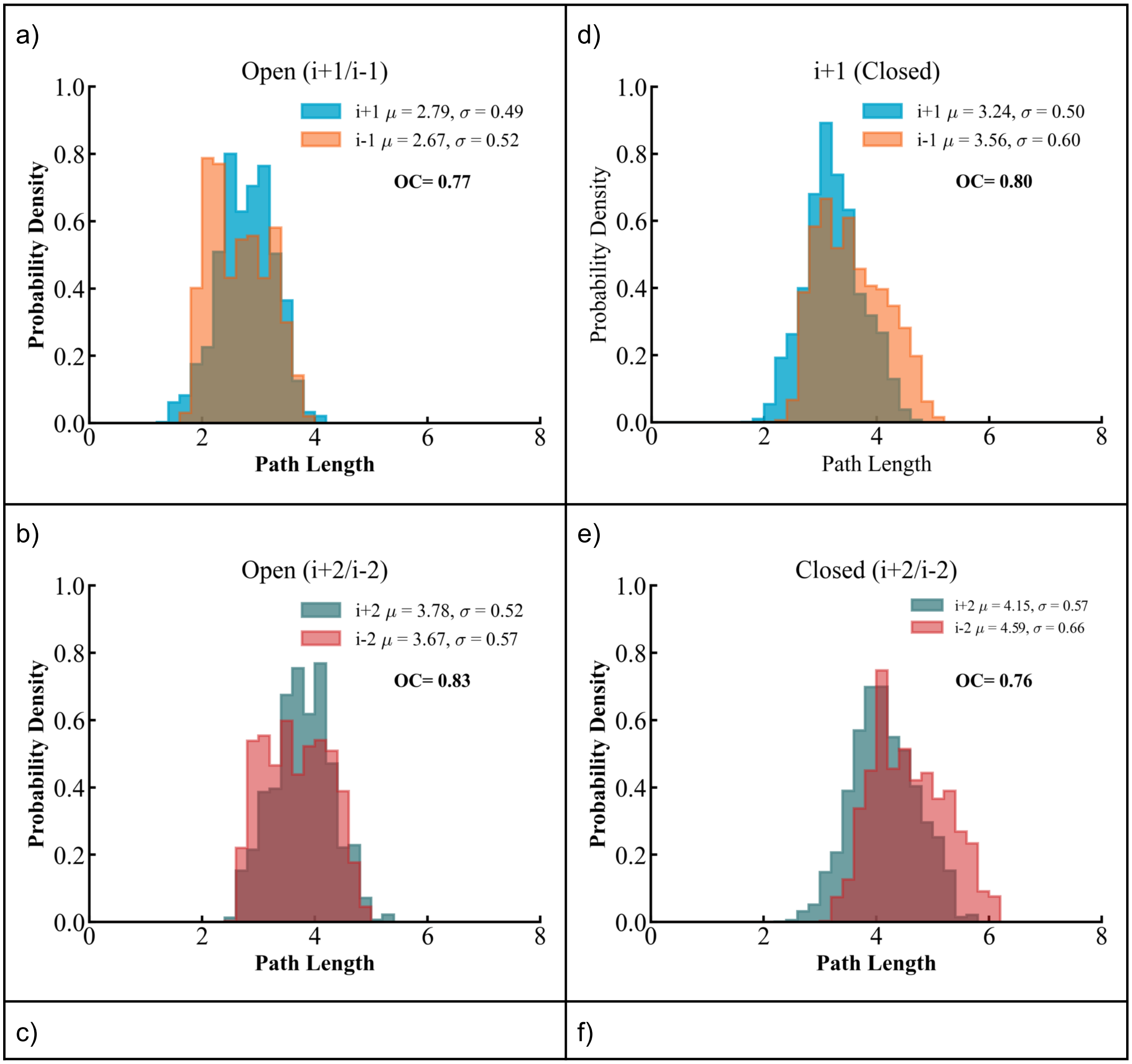
a)
Open (i+1/i-1)
i+1 $\mu$ = 2.79, $\sigma$ = 0.49
i-1 $\mu$ = 2.67, $\sigma$ = 0.52
OC= 0.77
Probability Density
Path Length
d)
i+1 (Closed)
i+1 $\mu$ = 3.24, $\sigma$ = 0.50
i-1 $\mu$ = 3.56, $\sigma$ = 0.60
OC= 0.80
Probability Density
Path Length
b)
Open (i+2/i-2)
i+2 $\mu$ = 3.78, $\sigma$ = 0.52
i-2 $\mu$ = 3.67, $\sigma$ = 0.57
OC= 0.83
Probability Density
Path Length
e)
Closed (i+2/i-2)
i+2 $\mu$ = 4.15, $\sigma$ = 0.57
i-2 $\mu$ = 4.59, $\sigma$ = 0.66
OC= 0.76
Probability Density
Path Length
c)
f)

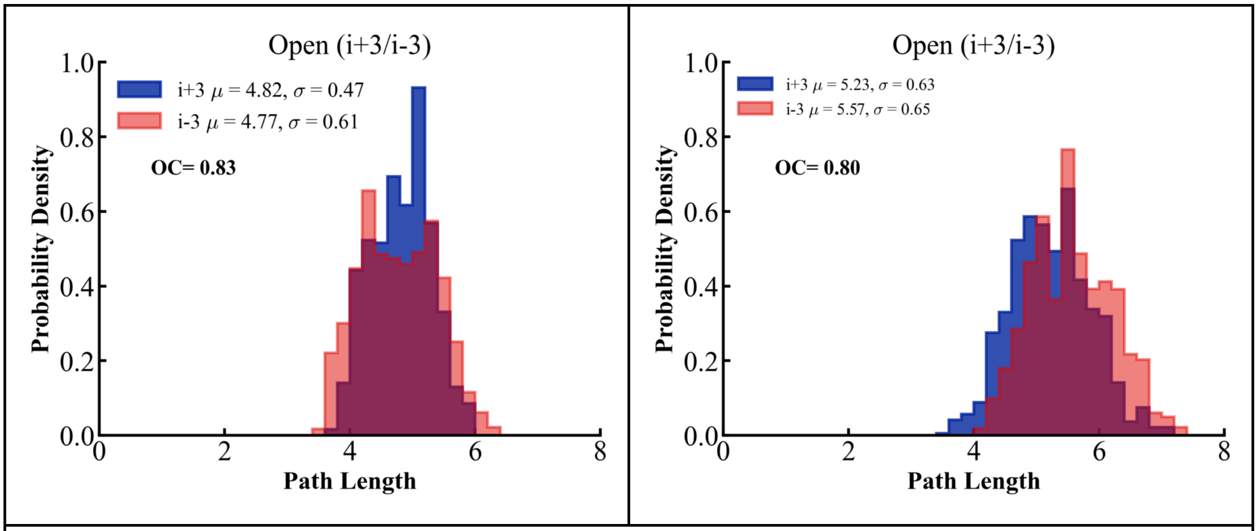


**FigureS7:** Path length distributions for equidistant protomer loops in ClpP configurations. (a)-(c) Open configuration, (d)-(f) Closed configuration
